# ATP-driven conformational dynamics reveal hidden intermediates in a heterodimeric ABC transporter

**DOI:** 10.64898/2026.02.12.705656

**Authors:** Matija Pečak, Christoph Nocker, Robert Tampé

## Abstract

ATP-binding cassette (ABC) transporters are essential molecular machines whose conformational dynamics have largely been inferred from ensemble-averaged measurements. Resolving dynamic heterogeneity and transient intermediates, however, requires single-molecule approaches. Here, we use single-molecule Förster resonance energy transfer (smFRET) to resolve ATP-driven conformational dynamics of the heterodimeric type IV ABC transporter TmrAB, a functional homolog of the human antigen transporter TAP, at the level of individual molecules. Fluorophores positioned at the nucleotide-binding domains and periplasmic gate were validated by accessible-volume simulations, fluorescence lifetimes, and ensemble FRET, demonstrating that these reporters reliably track conformational transitions. Single-molecule analysis distinguishes ATP-free and ATP-bound states and quantifies ATP-dependent population shifts from nucleotide-free to physiological ATP concentrations. Kinetic analysis further reveals an unexpectedly long ATP-bound dwell time of ∼300 ms. Using complementary stabilization strategies, we directly resolve a previously hidden outward-facing open state that is kinetically masked under turnover conditions. These results provide the first single-molecule characterization of TmrAB and establish a quantitative single-molecule framework for dissecting ATP-coupled conformational dynamics in heterodimeric ABC transporters.

**Impact Statement:** ATP-driven single-molecule imaging uncovers hidden outward-facing intermediates and unexpectedly long-lived ATP-bound states in the heterodimeric ABC transporter TmrAB, revealing how conformational heterogeneity shapes transport dynamics.

## Introduction

ATP-binding cassette (ABC) transporters constitute the largest family of primary active membrane transport systems, conserved across all domains of life^1–3^. Despite considerable structural diversity, all ABC transporters share a modular architecture comprising two conserved nucleotide-binding domains (NBDs)–the defining hallmark of the family–and two transmembrane domains (TMDs) that form the substrate translocation pathway^3,4^. Based on their TMD architecture, ABC transporters are classified into seven types that encompass importers, exporters, extractors, and mechanotransmitters^5^. Substrate translocation is driven by large conformational changes that are chemo-mechanically coupled to ATP binding, hydrolysis, and phosphate/ADP release^2,3^. ABC transporters play central roles in cellular homeostasis, nutrient uptake, waste removal, and toxin defense. Their dysfunction and misregulation are linked to numerous diseases and drug resistance^6^.

The heterodimeric type IV ABC transporter TmrAB from *Thermus thermophilus* has emerged as a powerful model system due to its exceptional thermal stability and functional homology to the transporter associated with antigen processing (TAP1/2), a key component of adaptive immunity^7–9^. Notably, TmrAB shares overlapping peptide specificity with TAP and can restore antigen presentation in TAP-deficient human cells^10^. Its inherent asymmetry, with one catalytically active (canonical) and one inactive (noncanonical) nucleotide-binding site (NBS), provides a unique opportunity to investigate functional specialization and asymmetry in ABC transport mechanisms.

Extensive structural studies, particularly using cryogenic electron microscopy (cryo-EM), have delineated the conformational landscape of TmrAB and yielded a detailed model of its translocation cycle^11,12^. In this model, TmrAB fluctuates between inward-facing wide and narrow conformations (IF^wide^ and IF^narrow^), characterized by a sealed periplasmic gate (PG) and well-separated NBDs, thereby permitting substrate access to the central binding cavity. ATP binding to both NBDs induces NBD dimerization and drives the transition into the outward-facing (OF) states, including an OF open (OF^open^) conformation with an open PG that enables substrate release into the periplasm, as well as an OF occluded (OF^occluded^) state characterized by a sealed PG and dimerized NBDs. Subsequent ATP hydrolysis and phosphate release lead to asymmetric unlocked return states (UR^asym^ and UR^asym^*), before the transporter returns to the IF conformation. These UR states feature a sealed PG, a partially open ADP-bound canonical NBS, and a tightly ATP-occluded noncanonical NBS^11^.

Single-turnover experiments have shown that ATP binding, rather than hydrolysis, drives the IF-to-OF transition, while phosphate release precedes the OF-to-IF switch^12–14^. Complementary ensemble approaches, including pulsed electron–electron double resonance (PELDOR/DEER) spectroscopy, have further characterized ATP-dependent conformational changes^15,16^. However, ensemble averaging inherently masks molecular heterogeneity, obscures inactive or misfolded subpopulations, and limits access to kinetic information.

Single-molecule techniques overcome these limitations by resolving conformational dynamics at the level of individual molecules^17,18^. In particular, single-molecule Förster resonance energy transfer (smFRET) enables real-time monitoring of protein conformational changes with nanometer precision^19–22^. Applied to ABC transporters, smFRET provides a unique opportunity to dissect transport cycles, resolve transient intermediates, and extract kinetic and mechanistic insights that remain inaccessible to ensemble-based measurement approaches^23–25^.

Here, we apply total internal reflection fluorescence (TIRF) microscopy combined with alternating laser excitation (ALEX)-based smFRET to detergent-solubilized heterodimeric ABC transporter TmrAB, providing the first single-molecule characterization of this system. By strategically positioning fluorophore pairs, we directly monitor ATP-dependent NBD dimerization and periplasmic gate (PG) opening, quantify conformational state occupancies across ATP concentrations ranging from nucleotide-free to physiological levels (3 mM), and uncover conformational dynamics previously masked by ensemble averaging. Using three orthogonal trapping strategies–(i) a slow-turnover mutant^11,12^, (ii) Mg²⁺ depletion^14,15^, and (iii) substrate trans-inhibition^26,27^–we resolved a previously hidden outward-facing open (OF^open^) state that rapidly exchanges with the outward-facing occluded (OF^occluded^) state. Distance measurements derived from smFRET closely matched predictions from accessible-volume (AV) simulations, cryo-EM structures, and PELDOR/DEER spectroscopy, confirming that detergent-solubilized TmrAB retains a native-like conformational landscape. Together, these results provide the first single-molecule quantification of conformational state occupancies for a heterodimeric type IV ABC transporter and establish TmrAB as a versatile model for single-molecule studies of ABC transport systems.

## Results

### Design of FRET-labeled TmrAB variants to probe conformational dynamics

To monitor conformational changes in distinct regions of TmrAB, we engineered double-cysteine variants for smFRET targeting the nucleotide-binding domains (NBDs) and the periplasmic gate (PG). The NBDs undergo ATP-dependent dimerization and post-hydrolysis dissociation, whereas the PG opening and closing controls substrate release into the periplasm^2,3,11^. Probing both regions provides complementary readouts of cytosolic and periplasmic coupling during transport.

Labeling positions were selected based on prior PELDOR/DEER studies^16^. The NBD reporter variant (TmrA^C416^B^L458C^, referred to as TmrAB^NBD^) monitors the noncanonical nucleotide-binding site (NBS), while the PG reporter (TmrA^C416A, T61C^B^R56C^, hereafter TmrAB^PG^) reports PG opening. In TmrAB^NBD^, the native single cysteine (C416) was retained for labeling, whereas in TmrAB^PG^ it was substituted by alanine to prevent off-target labeling. Using the noncanonical NBS prevents direct distinction between OF^occluded^ and asymmetric unlocked return states (UR^asym^ and UR^asym^*)^11^, reducing the number of resolvable FRET states and simplifying data interpretation.

Both variants were labeled with photostable LD555/LD655 fluorophores containing a 1,3,5,7-cyclooctatetraene moiety to suppress photobleaching and blinking^28,29^. Accessible-volume (AV) simulations^30^ across nine cryo-EM structures^11^ confirmed that donor–acceptor distances (*R*_DA_) and simulated FRET efficiencies (*E*_sim_) lie within the measurable range (**Fig. 1** and **Table 1**). For TmrAB^NBD^, *E*_sim_ shifts from 0.62 ± 0.02 (57.9 ± 0.7 Å, NBDs separated) to 0.84 ± 0.01 (45.2 ± 0.4 Å, NBDs dimerized), corresponding to predicted Δ*E*_sim_ of 0.22 ± 0.02 and Δ*R*_DA_ of 12.7 ± 0.8 Å. For TmrAB^PG^, *E*_sim_ changes from 0.96 ± 0.02 (30.6 ± 0.4 Å, closed PG) to 0.69 ± 0.03 (53.8 ± 2.0 Å, open PG), corresponding to predicted Δ*E*_sim_ of 0.27 ± 0.04 and Δ*R*_DA_ of 23.2 ± 2.0 Å.

**Figure 1.**
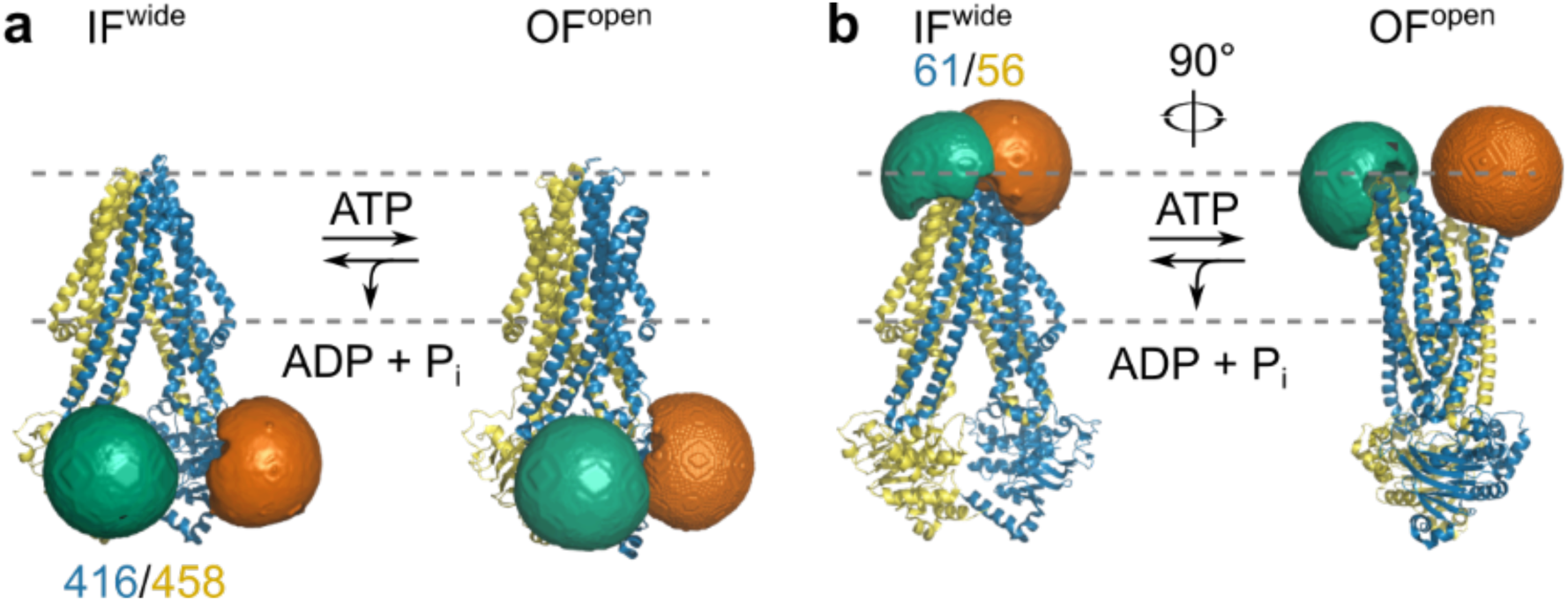
Accessible-volume (AV) simulations of LD fluorophores on TmrAB variants. AV simulations were performed for LD555 (donor) and LD655 (acceptor) fluorophores attached to the selected TmrAB labeling sites to assess whether donor–acceptor distances are suitable for smFRET measurements^30^. **a**, TmrAB^NBD^ (TmrA^C416^B^L458C^) and **b**, TmrAB^PG^ (TmrA^C416A, T61C^B^R56C^) in the inward-facing wide (IF^wide^; PDB: 6RAN, left) and outward-facing open (OF^open^; PDB: 6RAH, right) conformations. The approximate membrane position is indicated by the dashed grey line. TmrA is shown in blue with LD655 (orange) and TmrB in yellow with LD555 (green), with labeling positions indicated (416/458 for TmrAB^NBD^; 61/56 for TmrAB^PG^). For illustration, fluorophores are shown in a defined configuration; however, in experiments donor and acceptor dyes are stochastically attached, resulting in random distribution between subunits.

**Table 1:** Accessible-volume (AV) simulations of LD fluorophores on TmrAB variants. AV simulations were performed for LD555 (donor) and LD655 (acceptor) fluorophores attached to the selected TmrAB labeling sites to assess whether donor–acceptor distances are suitable for smFRET measurements^30^.AV simulations confirm that donor–acceptor distances (*R*_DA_) remain within the FRET-sensitive range in both conformations, predicting measurable changes in simulated FRET efficiencies (*E*_sim_). Cryo-EM structures of TmrAB reconstituted in lipid nanodiscs^11^ were used as templates, including apo and ATP-bound (3 mM ATP) states. Outward-facing open (OF^open^) and outward-facing occluded (OF^occluded^) structures were obtained via orthovanadate trapping (V_i_) or the slow-turnover mutant TmrA^E523Q^B (EQ).

| Conformation |  | PDB ID | TmrAB<sup>NBD</sup> |  | TmrAB<sup>PG</sup> |  |
| --- | --- | --- | --- | --- | --- | --- |
|  |  |  | RDA (Å) | Esim | RDA (Å) | Esim |
| IF<sup>narrow</sup> | (turnover) | 6RAF | 58.6 | 0.60 | 33.0 | 0.95 |
| IF<sup>wide</sup> | (turnover) | 6RAG | 57.2 | 0.63 | 30.2 | 0.96 |
| IF<sup>wide</sup> | (apo) | 6RAN | 57.8 | 0.62 | 30.3 | 0.96 |
| OF<sup>open</sup> | (V<sub>i</sub>) | 6RAJ | 44.8 | 0.84 | 55.8 | 0.66 |
| OF<sup>open</sup> | (EQ) | 6RAH | 44.9 | 0.84 | 51.8 | 0.72 |
| OF<sup>occluded</sup> | (V<sub>i</sub>) | 6RAK | 45.4 | 0.83 | 27.0 | 0.98 |
| OF<sup>occluded</sup> | (EQ) | 6RAI | 45.5 | 0.83 | 32.4 | 0.95 |
| UR<sup>asym</sup> | (turnover) | 6RAL | 45.9 | 0.83 | 31.2 | 0.96 |
| UR<sup>asym*</sup> | (turnover) | 6RAM | 47.6 | 0.80 | 31.2 | 0.96 |

A slow-turnover TmrAB variant (TmrA^C416A, E523Q, T61C^B^R56C^, hereafter TmrAB^PG_EQ^) was also generated to kinetically stabilize ATP-bound conformations. This mutation reduces ATP hydrolysis by ∼1000-fold, extending the catalytic half-life of ∼25 min at 45 °C^11,12,14,15^.

### TmrAB variants are suitable for FRET studies

TmrAB variants were expressed in *E. coli* and purified by immobilized metal-affinity chromatography. SDS-PAGE and size-exclusion chromatography (SEC) confirmed purity and monodispersity of the samples (**Fig. 1–Fig. S1a,b**). ATPase assays showed that TmrAB^wt^ retained enzymatic activity after purification, yielding a Michaelis-Menten constant (*K*_m_) of 0.97 ± 0.28 mM and turnover rate (*k*_cat_) of 2.57 ± 0.38 s^-1^ at 40 °C (**Fig. 1–Fig. S1c**), consistent with previous reports^9,10,12^.

Site-specific labeling yielded >90% labeling efficiency per cysteine, with ∼40–50% per-site labeling efficiency for donor-only and acceptor-only populations (**Fig. 1–Fig. S1d–f**). Stochastic labeling of the TmrAB variants results in both homo (donor-donor, acceptor-acceptor) and hetero (donor-acceptor) labeled species. Only complexes with appropriate donor–acceptor stoichiometry were included in subsequent analyses.

Fluorescence lifetime (*τ*) analysis confirmed that the conjugated fluorophores retained their photophysical properties and sufficient rotational freedom for reliable FRET measurements. *τ* histograms of both conjugated and free fluorophores were fitted using a biexponential decay model, from which amplitude-weighted average lifetimes were calculated. For TmrAB^NBD^, average *τ* values were 0.93 ± 0.02 ns (LD555) and 1.52 ± 0.01 ns (LD655), whereas TmrAB^PG^ exhibited average *τ* values of 0.95 ± 0.02 ns (LD555) and 1.65 ± 0.01 ns (LD655) (**Fig. 1–Fig. S2a**). By comparison, free dyes in buffer displayed lifetimes of 1.12 ± 0.02 ns (LD555) and 1.29 ± 0.01 ns (LD655). The similarity between free and conjugated dye lifetimes, together with comparable values across variants, indicates minimal protein-induced quenching and supports sufficient orientational averaging for quantitative smFRET analysis^18,31–34^.

Ensemble ATP titration (0–10 mM ATP) showed ATP-dependent donor quenching and acceptor sensitization (**Fig. 1–Fig. S2b–d**). ATP-induced fluorescence changes provided a quantitative readout of conformational transitions and enabled estimation of apparent equilibrium dissociation constants for ATP binding (*K*_d, ATP_) in labeled TmrAB variants (**Fig. 1–Fig. S2e–g**). The measured apparent *K*_d, ATP_ values were 51 ± 38 μM for TmrAB^NBD^, 68 ± 25 μM for TmrAB^PG^, and 95 ± 26 μM for the slow-turnover variant TmrAB^PG_EQ^, consistent with prior biochemical measurements (∼100 µM for TmrA^E523Q^B)^12^. Labeling did not measurably perturb ATP binding.

Previous studies on spin-labeled TmrAB^NBD^ demonstrated transport activity comparable to wild-type TmrAB^16^, while AV simulations confirmed that fluorophores at these positions do not interfere with ATP- or substrate-binding sites. For labeled TmrAB^PG^ variant, no previous transport activity had been reported; we therefore performed transport assays (**Fig. 1**–**Fig. S3**) which confirmed that LD555/LD655 labeling at the PG does not impair function, as labeled and wild-type TmrAB showed indistinguishable substrate transport in single-liposome assays (**Fig. 1**–**Fig. S3c**).

### ATP-induced conformational switching resolved by smFRET

TmrAB was immobilized on PEGylated surfaces using a conformation-independent, TmrB-specific nanobody (Nb9F10^S63C^)^11^. Previous studies confirmed that Nb9F10^S63C^ does not perturb TmrAB transport^11,14^, consistent with single-liposome transport assays performed in this study (**Fig. 1–Fig. S3d**). For surface immobilization, Nb9F10^S63C^ was conjugated to maleimide-PEG_11_-biotin, serving as an anchor to attach TmrAB to surface-immobilized streptavidin (**Fig. 2a**). Donor and acceptor photons were recorded by a total-internal reflection fluorescence (TIRF) microscope using alternating laser excitation (ALEX; NanoImager) at 40 °C (**Fig. 2b**). Single-molecule localization, fluorescence-trajectory extraction, and background correction were performed using NanoImager software, followed by DeepFRET-based machine-learning trace classification and corrections for donor leakage, direct acceptor excitation, and differences in detection-efficiency^18,35,36^ (**Fig. 2–Fig. S1** and **Fig. 2–Fig. S2)**. FRET efficiency (*E*) and stoichiometry (*S*) were calculated from the corrected fluorescence trajectories (see **Methods, Eq. 1** and **Eq. 2**), and 2D *E* versus *S* diagrams were constructed to verify that only complexes with appropriate donor–acceptor stoichiometry (*S* ≈ 0.5) were included in the smFRET analysis (**Fig. 2c**).

**Figure 2.**
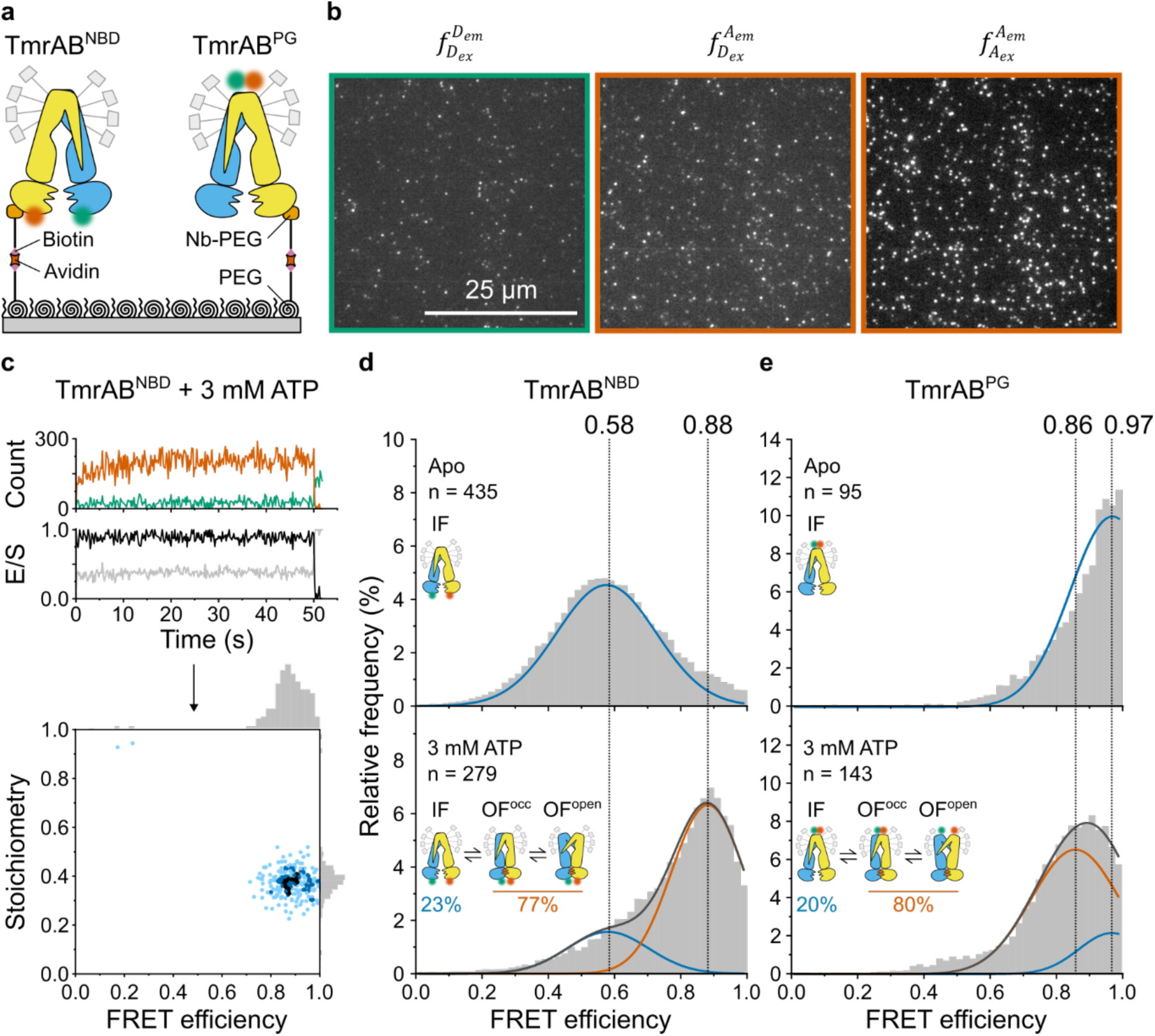
ATP-induced conformational changes of TmrAB analyzed by smFRET. **a**, Experimental setup. TmrAB^NBD^ (left) and TmrAB^PG^ (right) were labeled with LD555/LD655 and immobilized on PEGylated coverslips via a biotinylated conformation-independent, TmrB-specific nanobody (Nb9F10^S63C^)^11,14^. **b**, smFRET imaging was performed using total internal reflection fluorescence (TIRF) microscopy with alternating laser excitation (ALEX; donor: 532 nm; acceptor: 640 nm). Emission was collected in donor (498-620 nm) and acceptor (662-710 nm) channels. Donor emission upon donor excitation 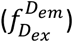, sensitized acceptor emission upon donor excitation 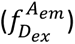, and direct acceptor emission upon acceptor excitation 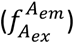 were used to calculate FRET efficiency (*E*) and stoichiometry (*S*). Data were analyzed using DeepFRET^35^. **c**, Representative TmrAB^NBD^ trace in ATP-bound state (top) and corresponding E/S plot (bottom). Donor emission upon donor excitation is shown in green (photon counts per frame), acceptor emission upon donor excitation in orange, FRET efficiency (*E*) in black, and stoichiometry (*S*) in grey. The initial increase in donor fluorescence reflects photophysical equilibration and instrumental stabilization and does not affect E or S, which are ratio-based and time-invariant^18^. **d**,**e**, Population analysis. FRET efficiency (*E*) histograms for (**d**) TmrAB^NBD^ and (**e**) TmrAB^PG^ are shown for apo (top) and ATP-bound (bottom; 3 mM ATP) conditions. Histograms were fitted with two Gaussian populations corresponding to the apo state (blue; defined from apo measurements) and ATP-bound state (orange; defined from saturating ATP fits). Dotted vertical lines indicate mean *E* values of each population, and relative fractions (Gaussian areas) are summarized schematically in each panel.

FRET efficiency (*E*) histograms revealed two dominant Gaussian populations corresponding to apo and ATP-bound states (**Fig. 2d,e**). Low-amplitude features were excluded from quantitative analysis because they were not reproducibly observed across replicates, lacked structural support from cryo-EM and DEER/PELDOR data, introduced poorly constrained fitting parameters, and increased the risk of overfitting. In the absence of ATP, only the apo population was observed, whereas addition of 3 mM ATP induced the appearance of a second ATP-bound population. For TmrAB^NBD^, the apo and ATP-bound populations exhibited mean *E* values of 0.58 and 0.88 (Δ*E* = 0.30), respectively, with ∼77% of molecules occupying the ATP-bound state. For TmrAB^PG^, mean *E* values were 0.97 (apo) and 0.86 (ATP-bound) (Δ*E* = 0.11), with ∼80% of molecules in the ATP-bound state.

Distance estimates using a Förster radius of *R*_0_ = 63.5 Å (ref.^37^) yielded apparent distances of 60.2 Å (apo) and 45.6 Å (ATP-bound) for TmrAB^NBD^, and 35.6 Å (apo) and 46.9 Å (ATP-bound) for TmrAB^PG^. For TmrAB^NBD^, the experimentally derived Δ*R* of 14.6 Å closely agreed with the AV simulations. In contrast, the smaller Δ*R* of 11.4 Å observed for TmrAB^PG^ deviated from simulated values, suggesting that the ATP-bound population at this site represents a mixture of rapidly interconverting states rather than a single well-defined state.

### ATP dependence of conformational equilibria

To assess the ATP sensitivity, we quantified conformational responses across a wide range of ATP concentrations, spanning well below the reported *K*_d, ATP_ (∼100 µM for TmrA^E523Q^B)^12^ up to physiologically relevant levels (3 mM ATP). ATP concentrations up to 3 mM were selected to approximate near-physiological conditions commonly used in in-vitro studies, where millimolar ATP levels ensure saturation of nucleotide-dependent conformational transitions and facilitate comparison with previous biochemical and structural analyses^10–12,14^. smFRET measurements revealed dose-dependent population shifts: TmrAB^NBD^ shifted from a low-FRET apo state (*E* = 0.58) to a high-FRET ATP-bound state (*E* = 0.88), whereas TmrAB^PG^ shifted from a high-FRET apo state (*E* = 0.97) to a lower-FRET ATP-bound state (*E* = 0.86) (**Fig. 3a,c**). Langmuir isotherm fits yielded apparent *K*_d, ATP_ values of 13 ± 1 μM for TmrAB^NBD^ and 2 ± 1 μM for TmrAB^PG^ (**Fig. 3b,d**), significantly lower than *K*_d ATP_ determined in ensemble FRET measurements (**Fig. 1–Fig. S2e–g)**. This difference likely arises from the difficulty of deconvoluting overlapping FRET populations at sub-*K*_d, ATP_ concentrations, particularly for TmrAB^PG^, where state assignment is less well separated. Despite this quantitative offset, both approaches consistently indicate ATP saturation well below physiological concentrations and therefore support the same mechanistic conclusion that ATP binding drives conformational switching in TmrAB.

**Figure 3.**
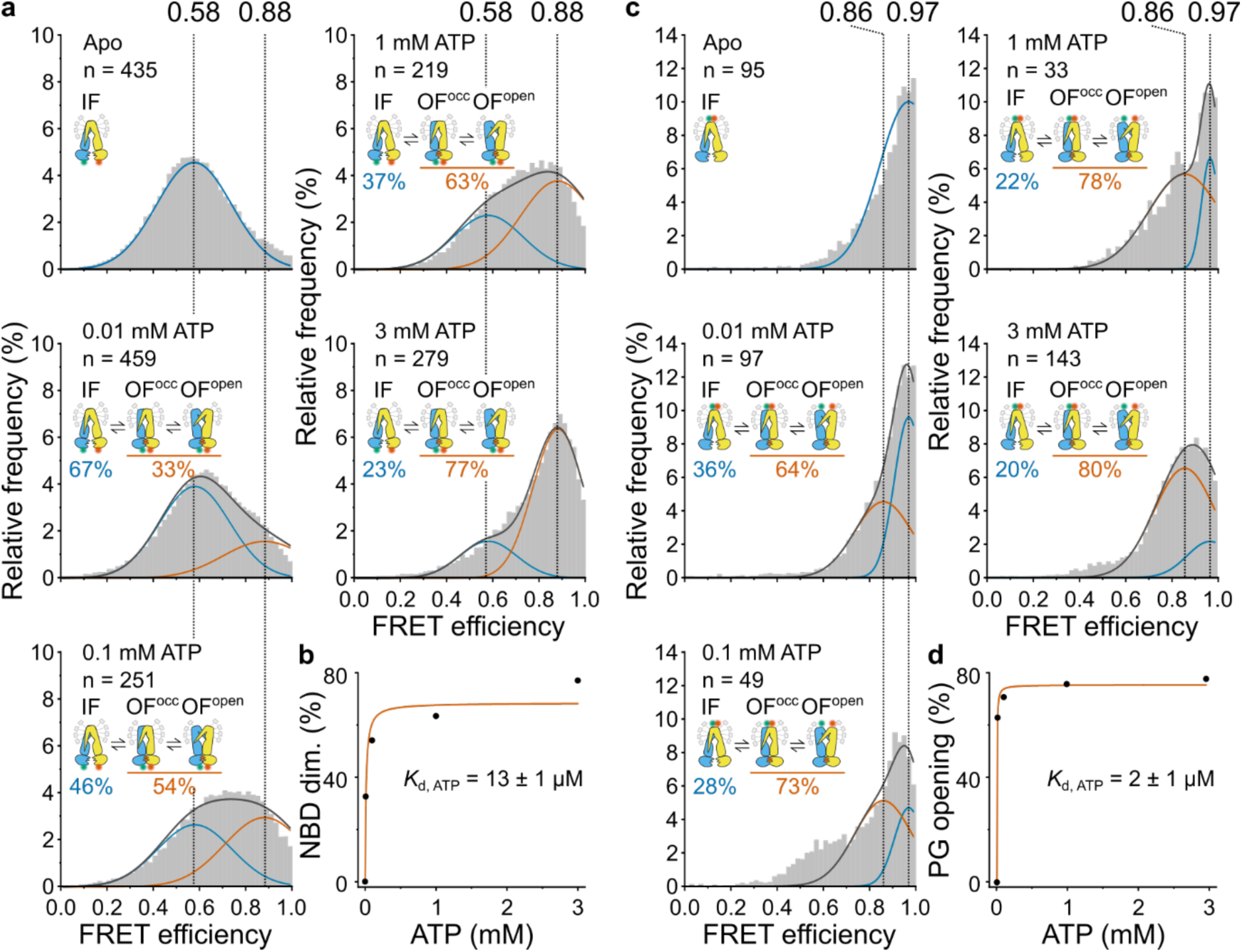
ATP-dependent shifts in smFRET populations of TmrAB. **a**,**c**, Increasing ATP concentrations (0-3 mM) progressively shifted the population between apo and ATP-bound conformations for (**a**) TmrAB^NBD^ and (**c**) TmrAB^PG^. FRET efficiency (*E*) histograms were fitted with two Gaussian populations corresponding to the ATP-free state (blue; defined from apo samples) and the ATP-bound state (orange; determined from saturating ATP conditions). Dotted vertical lines indicate the mean *E* values of each state, and relative fractions (Gaussian areas) are summarized schematically in each panel. **b**,**d**, ATP-binding curves obtained by plotting the fraction of molecules in the ATP-bound states as a function of ATP concentration for (**b**) TmrAB^NBD^ (reporting NBD dimerization) and (**d**) TmrAB^PG^ (reporting PG opening). Data were fitted with a Langmuir isotherm to determine the apparent dissociation constant *K*_d, ATP_ of each variant.

### Trapping of TmrAB^PG^ reveals a previously hidden outward-facing open state

At 3 mM, ∼80% of TmrAB^PG^ complexes populated the ATP-bound state (*E* = 0.86), closely matching the ATP-bound population (∼77% NBD-dimerized) observed for TmrAB^NBD^ (**Fig. 3**). This agreement indicates that both labeling strategies consistently report ATP-dependent conformational changes. However, the ATP-induced shift observed for the periplasmic-gate reporter TmrAB^PG^ (Δ*E* = 0.11, Δ*R* = 11.4 Å; **Fig. 3c**) was substantially smaller than predicted by AV simulations (Δ*E* = 0.27, Δ*R* = 23.2 Å; **Table 1**). This discrepancy suggests that the ATP-bound population at *E* = 0.86 represents an unresolved ensemble, potentially comprising OF^open^ and OF^occluded^ conformations as well as the post-hydrolysis asymmetric unlocked return states (UR^asym^ and UR^asym^*), which cannot be distinguished from OF^occluded^ within the current FRET geometry^11^.

To determine whether these states are kinetically unresolved by smFRET, we applied three complementary strategies to arrest the OF^open^ conformation of TmrAB^PG^: (i) a slow-turnover mutant (TmrAB^PG_EQ^), (ii) Mg²⁺ depletion using EDTA, and (iii) trans-inhibition by high concentrations of the substrate peptide R9L (RRYQKSTEL) (**Fig. 4**). Slow-turnover variants have previously enabled structural separation of OF^open^ and OF^occluded^ states^11,12,14,15^. Mg²⁺ depletion blocks ATP hydrolysis while preserving ATP binding, thereby enabling rapid and reversible trapping^14^. We further hypothesized that trans-inhibition by peptide binding sterically restricts PG closure and would therefore stabilize OF^open^ in a dose-dependent manner^26,27^.

**Figure 4.**
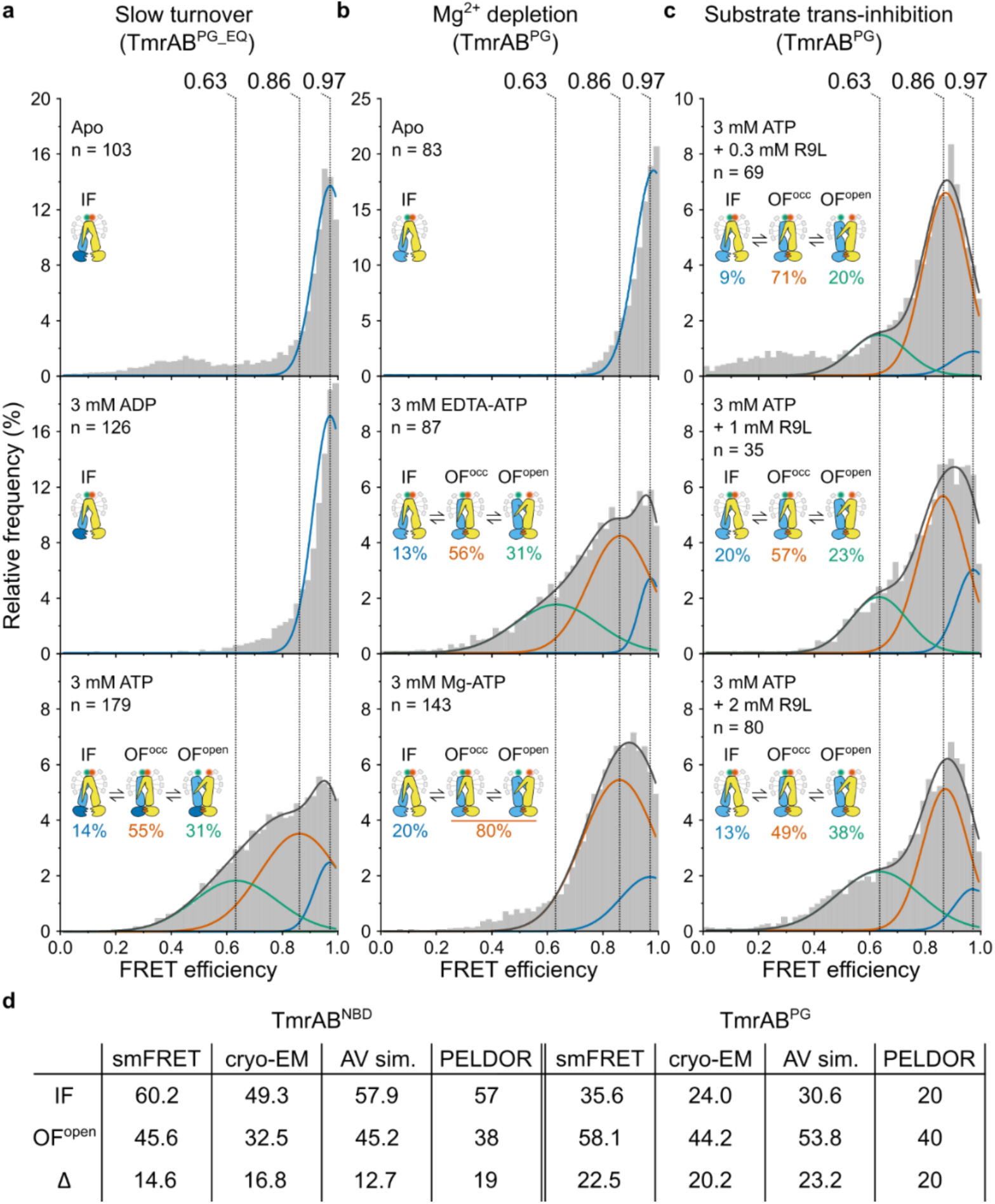
Identification of the outward-facing open (OF^open^) conformation. **a–c**, Three complementary approaches were used to resolve the OF^open^ state: (**a**) the slow-turnover variant TmrAB^PG_EQ^, (**b**) imaging in Mg^2+^-free buffer supplemented with EDTA, and (**c**) stabilizing via trans-inhibition using high concentrations of peptide substrate R9L (0.3–2 mM). FRET efficiency (*E*) histograms were fitted with three Gaussian populations corresponding to the ATP-free state (blue), the ATP-bound state (orange), and the OF^open^ state (green). All three strategies reveal a distinct OF^open^ population. Dotted vertical lines indicate the mean *E* values of each state, and relative fractions (Gaussian areas) are summarized schematically in each panel. **d**, Comparison of inter-residue distances. Distances (Å) between selected residues on the NBDs and PG of TmrAB (C_β_–C_β_) were determined using smFRET (this study, detergent-solubilized TmrAB), cryo-EM structures of nanodisc-reconstituted TmrAB (PDB 6RAH, 6RAN)^11^; accessible-volume (AV) simulations (this study), and PELDOR/DEER measurements of detergent-solubilized TmrAB^16^.

In the absence of ATP, either without nucleotide or in the presence of ADP (3 mM), the slow-turnover variant TmrAB^PG_EQ^ populated a single high-FRET state (*E* = 0.97) (**Fig. 4a**, top and middle). These results indicate that ADP binding alone is insufficient to promote either NBD dimerization or PG opening, which is consistent with previous biochemical and structural observations^11,12^. Upon ATP addition (3 mM), however, the conformational landscape diverged sharply from that of TmrAB^PG^. Instead of the two-state distribution observed for TmrAB^PG^ (apo: *E* = 0.97, ∼20%; ATP-bound: *E* = 0.86, ∼80%) (**Fig. 2e**, bottom), TmrAB^PG_EQ^ exhibited three well-resolved populations with mean *E* values of 0.97 (∼14%), *E* = 0.86 (∼55%), and *E* = 0.63 (∼31%) (**Fig. 4a**, bottom).

As in TmrAB^PG^, the high-FRET population (*E* = 0.97) corresponds to the IF state, whereas the intermediate-FRET population (*E* = 0.86) likely represents a dynamic equilibrium between OF^open^ and OF^occluded^ conformations, as suggested by cryo-EM analyses^11^. Post-hydrolysis return states (UR^asym^ and UR^asym^*) are expected to be minimally populated in TmrAB^PG_EQ^ because of its drastically reduced ATP hydrolysis rate^12^. Notably, the low-FRET ATP-bound population (*E* = 0.63) was entirely absent in wild-type TmrAB^PG^. The transition from *E* = 0.97 to *E* = 0.63 corresponds to Δ*E* = 0.34 and Δ*R* = 22.5 Å, in close agreement with the IF→OF^open^ distance predicted by AV simulations (Δ*R* = 23.2 Å; **Fig. 1** and **Table 1**), thereby supporting assignment of the *E* = 0.63 population to the OF^open^ conformation.

Mg²⁺ depletion independently reproduced this three-state landscape. In the absence of Mg^2+^, wild-type TmrAB^PG^ transitioned from a single apo population (without ATP; *E* = 0.97) (**Fig. 4b**, top) to three ATP-bound populations (3 mM ATP; *E* = 0.97, ∼13%; *E* = 0.86, ∼56%; *E* = 0.63, ∼31%) (**Fig. 4b**, middle), closely resembling those observed for the slow-turnover variant (**Fig. 4a**, bottom). Reintroducing Mg²⁺ abolished the *E* = 0.63 population and restored the wild-type TmrAB^PG^ two-state distribution (**Fig. 4b**, bottom). This reversibility confirmed that the ATP-bound OF^open^ state (*E* = 0.63) is selectively revealed only when ATP hydrolysis is prevented.

Finally, we tested whether periplasmic substrate binding shifts the conformational equilibrium of wild-type TmrAB^PG^ toward OF^open^. In the presence of ATP (3 mM), increasing concentrations of peptide substrate (0.3–2 mM R9L) progressively enriched the *E* = 0.63 population from ∼20% to ∼38% (**Fig. 4c**). This dose-dependent stabilization mirrors the trans-inhibition behavior reported for human TAP1/2 and reflects the upper substrate-loading capacity of the transporter^26^.

Distance changes derived from smFRET closely match AV simulations, cryo-EM structures (PDB 6RAH, 6RAN)^11^, and DEER/PELDOR measurements^15^ (**Fig. 4d**), together validating assignment of the *E* = 0.63 population as the OF^open^ conformation.

### Kinetics and thermodynamics of the transport cycle

ALEX-smFRET data were acquired with an effective temporal resolution of 200 ms (100 ms per excitation channel). Shorter integration times compromised the signal-to-noise ratio and precluded reliable FRET determination. To quantify conformational dynamics, we applied Hidden Markov Modeling (HMM) using MASH-FRET^38^, classifying traces as either static (single FRET state) or dynamic (multiple states). Approximately 95% of traces in each condition were classified as static. The predominance of static traces is consistent with most conformational transitions occurring faster than the 200 ms observation window rather than reflecting long-lived conformational arrest.

Although individual transitions could not be directly resolved, population-based analysis (**Fig. 3**), combined with biochemical turnover measurements (**Fig. 1–Fig. S1c**), allowed estimation of ATP-bound dwell times. Under saturating ATP conditions well above the apparent *K*_d, ATP_ (3 mM, 40 °C), wild-type TmrAB exhibited a catalytic turnover rate of *k*_cat_ = 2.57 ± 0.38 s^-1^ (**Fig. 1–Fig. S1c**), corresponding to a full transport cycle time (*τ*_cycle_) of 395 ± 55 ms. ATP-bound dwell times (*τ*_d_) were derived from population ratios obtained from Gaussian fits of the FRET efficiency histograms (**Fig. 3**; see Methods, Eq. 3 and Eq. 4). These analyses yielded ATP-bound dwell times of 304 ± 43 ms for TmrAB^NBD^ (∼77% ATP-bound) and 316 ± 44 ms for TmrAB^PG^ (∼80% ATP-bound), with the remaining ATP-free intervals (∼20–23%) accounting for 91 ms and 79 ms of the cycle, respectively.

The kinetic behavior of the ATP-bound ensemble is consistent with the thermodynamic properties previously determined for TmrAB. Although ATP-bound conformations accounted for most of the transport cycle (∼77–80% occupancy), individual OF^open^ and OF^occluded^ states could not be resolved within the ATP-bound period (∼310 ms), indicating rapid interconversion between these conformations. This observation agrees with previous thermodynamic analyses showing that ATP binding drives the IF-to-OF transition with an overall free-energy change close to zero (Δ*G* = 0.26 ± 0.74 kJ mol⁻¹), despite substantial compensating enthalpic and entropic contributions (Δ*H* = 30.14 ± 2.80 kJ mol⁻¹; *T*Δ*S* = 28.26 kJ mol⁻¹) (ref.^15^). This entropy–enthalpy compensation places the conformational equilibrium near its midpoint, minimizing the energetic bias between ATP-bound conformations and thereby promoting rapid exchange between OF^open^ and OF^occluded^ states. Accordingly, the unresolved ATP-bound ensemble observed during steady-state turnover is consistent with a thermodynamic landscape that enables dynamic sampling of multiple outward-facing conformations.

Together, these measurements establish a quantitative single-molecule description of both the kinetic progression and thermodynamic state occupancies throughout the catalytic cycle of a heterodimeric ABC transporter under active turnover conditions. By integrating population analysis with biochemical turnover measurements, this framework provides complementary insight into the dynamic and energetic landscape that governs the translocation cycle.

## Discussion

smFRET has become an indispensable tool for dissecting conformational dynamics of membrane proteins, including receptors, ion channels, and transporters, by directly linking structural transitions to functional states. With the ABC transporter family, however, smFRET studies have largely focused on monomeric or homodimeric systems^23–25^, leaving asymmetric heterodimeric transporters comparatively underexplored. Here, we extend single-molecule analysis to the heterodimeric type IV ABC transporter TmrAB, revealing dynamic features of the transport cycle that are inaccessible to ensemble-averaged approaches.

By positioning FRET reporters at the nucleotide-binding domains (NBDs) and periplasmic gate (PG), we directly monitored ATP-dependent coupling between chemical energy input and global conformational rearrangements. Importantly, NBD dimerization is not uniquely coupled to a single outward-facing state but can lead to either an outward-facing open (OF^open^) or occluded (OF^occluded^) conformation^11,12^. This decoupling highlights the need to monitor both cytosolic and periplasmic regions to resolve the transport mechanism.

A limitation of the present study is the use of two FRET reporter pairs. While these positions were selected to probe key functional regions and validated by structural and spectroscopic benchmarks, additional labeling positions could in principle resolve further intermediates. However, increasing the number of states would also complicate quantitative analysis due to partially overlapping FRET populations, particularly for TmrAB^PG^. The current design therefore balances mechanistic resolution with robust state assignment.

Labeling sites previously validated for PELDOR/DEER spectroscopy^10,15,16^ were adapted for smFRET and rigorously benchmarked using accessible-volume (AV) simulations^30^, fluorescence lifetime analysis, and ensemble FRET titrations. This validation is essential because fluorophores impose stricter steric and rotational constraints than spin labels^32^. Collectively, these controls demonstrate that fluorophore attachment preserves native-like conformational behavior and ATP responsiveness. Fluorescence lifetime measurements further indicated efficient energy transfer without evidence of substantial protein-fluorophore quenching or restricted mobility^18^.

Single-molecule measurements resolved two dominant FRET populations corresponding to apo and ATP-bound states for both TmrAB^NBD^ and TmrAB^PG^. ATP titrations from sub-*K*_d, ATP_ to physiological concentrations (3 mM ATP) produced gradual, concentration-dependent redistribution between these states, demonstrating high sensitivity of both reporters. Apparent *K*_d, ATP_ values derived from smFRET (2–13 µM) were lower than those obtained from ensemble FRET (50–100 µM). Importantly, both approaches agree that TmrAB reaches conformational saturation well below physiological ATP concentrations and retains Michaelis–Menten behavior.

Using three independent strategies—slow-turnover catalysis, Mg²⁺ depletion, and substrate trans-inhibition—we resolved this previously hidden OF^open^ conformation. The observed distance changes agree with cryo-EM structures^11^, PELDOR/DEER data^10,15,16^, and AV-based predictions, indicating that detergent-solubilized TmrAB samples a largely native-like conformational landscape.

Although detergent and lipid nanodisc environments yield similar global states, fluorophore behavior may still be influenced by membrane proximity in lipid systems^20,39,40^. AV simulations suggest that dyes near the periplasmic gate may experience steric constraints due to partial membrane overlap. While negligible under detergent conditions, this effect should be considered in future studies of membrane-reconstituted systems.

Substrate addition shifted the conformational equilibrium in a concentration-dependent manner, progressively stabilizing the OF^open^ state. This behavior is consistent with trans-inhibition observed in other ABC transporters, including TAP1/2^26^ and ABCC1^27^, and reflects finite substrate loading capacity^26^.

Quantitative deconvolution of FRET populations enabled estimation of conformational state occupancies under physiological ATP concentrations. During active turnover, TmrAB populates the IF state (∼20%), OF^open^ (∼25%), and OF^occluded^/post-hydrolysis states (UR^asym^ and UR^asym^*) (∼55%; **Fig. 5**). The current labeling scheme does not allow direct separation of OF^occluded^ from post-hydrolysis states due to fluorophore placement at the noncanonical NBS. However, post-hydrolysis contributions are expected to be reduced in the slow-turnover mutant TmrAB^PG_EQ^. To our knowledge, this represents the first single-molecule quantification of conformational equilibria in a heterodimeric ABC transporter under catalytic conditions. While IF substructure resolution is lower than in cryo-EM classifications^11,14^, smFRET uniquely captures ATP-bound intermediates that interconvert too rapidly for structural resolution.

**Figure 5.**
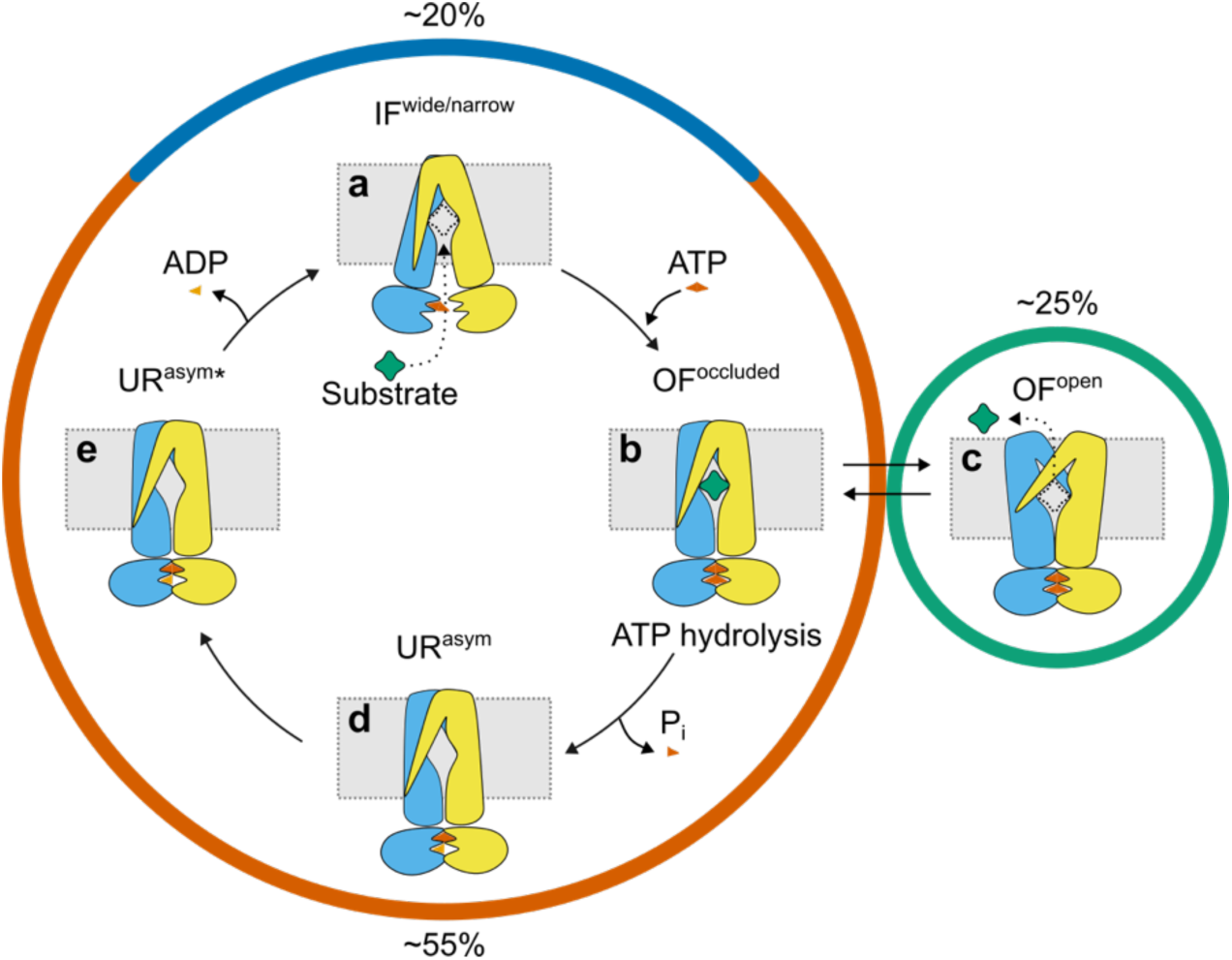
Conformational state distribution and catalytic cycle of TmrAB under active turnover. Schematic of the TmrAB transport cycle summarizing major conformational states and their estimated population distributions under physiological ATP concentrations (3 mM, 40 °C). **a**, The inward-facing apo state (IF^narrow^ and IF^wide^; blue arc) accounts for ∼20% of molecules and is characterized by separated NBDs and a cytosol-accessible substrate-binding cavity. Substrate binding stabilizes the IF^wide^ conformation^11^. ATP binding induces NBD dimerization and formation of the ATP-bound ensemble. **b**,**c**, Under substrate-bound turnover conditions, TmrAB proceeds via (**b**) OF^occluded^ and (**c**) OF^open^ states in which substrate release occurs. In steady-state turnover, the ATP-bound ensemble rapidly interconverts between OF^occluded^ and OF^open^, accounting for ∼25% of the ATP-bound population (green circle). These transitions occur faster than the ∼200 ms temporal resolution of smFRET measurements, resulting in an averaged signal under turnover conditions. OF^occluded^ is proposed as an obligate intermediate between IF and OF^open^, preventing substrate backflow by maintaining an occluded binding cavity during rearrangements of the PG and NBDs. Although a substrate-bound OF^occluded^ state has not been directly observed for TmrAB, its existence is supported by structures of the homodimeric type IV transporter BmrA^47^. Reduced ATP hydrolysis or substrate trans-inhibition enables trapping of the OF^open.^ state. **d**,**e**, ATP hydrolysis and phosphate (P_i_) release generate post-hydrolysis return states (**d**) UR^asym^ and (**e**) UR^asym^*. Subsequent ADP release restores the apo IF conformation, completing the transport cycle. Overall, the ATP-bound phase (**b–e**) represents ∼55% occupancy (orange arc) with an estimated dwell time of ∼310 ms, whereas the apo/ATP-rebinding phase (**a**) lasts ∼90 ms, yielding a total cycle time of ∼400 ms (*k*_cat_ = 2.57 s⁻¹). TmrA is shown in blue, TmrB in yellow, substrate as a green diamond, and nucleotides as orange symbols. Dotted grey boxes indicate the approximate position of the NBD dimer interface.

All experiments were performed at 40 °C for consistency with prior biochemical work and fluorophore stability. At the physiological temperature of *T. thermophilus* (68 °C), absolute rates are expected to increase, although relative state distributions may be preserved if the free-energy landscape remains unchanged. Despite a substantial fraction of static trajectories, ATP-dependent shifts indicate that TmrAB operates near the temporal resolution of the measurements. Combining state occupancies with enzymatic turnover yields an ATP-bound dwell time of ∼300 ms, consistent with previous biochemical estimates^9,12^.

Emerging microsecond-resolution smFRET approaches could directly resolve short-lived intermediates^41^, and multi-color FRET strategies^42,43^ may enable simultaneous monitoring of NBDs, PG, and nucleotide sites. These advances would allow direct detection of the OF^occluded^ states and post-hydrolysis dynamics.

Overall, the consistency between reporter positions, ensemble measurements, and structural data support the conclusion that the observed conformational states reflect native TmrAB behavior. While certain intermediates remain kinetically unresolved under turnover conditions, they are strongly supported by independent trapping approaches, reinforcing the robustness of the mechanistic model. Additional conditions, including nucleotide-state asymmetry and substrate-only binding, represent promising directions for future work.

In summary, this work establishes smFRET as a powerful approach for mapping the dynamic landscape of asymmetric ABC transporters. By quantitatively linking ATP binding, conformational equilibria, and kinetics at the single-molecule level, we define how chemical energy is converted into directional transport in heterodimeric ABC systems.

## Methods

### Key resources table

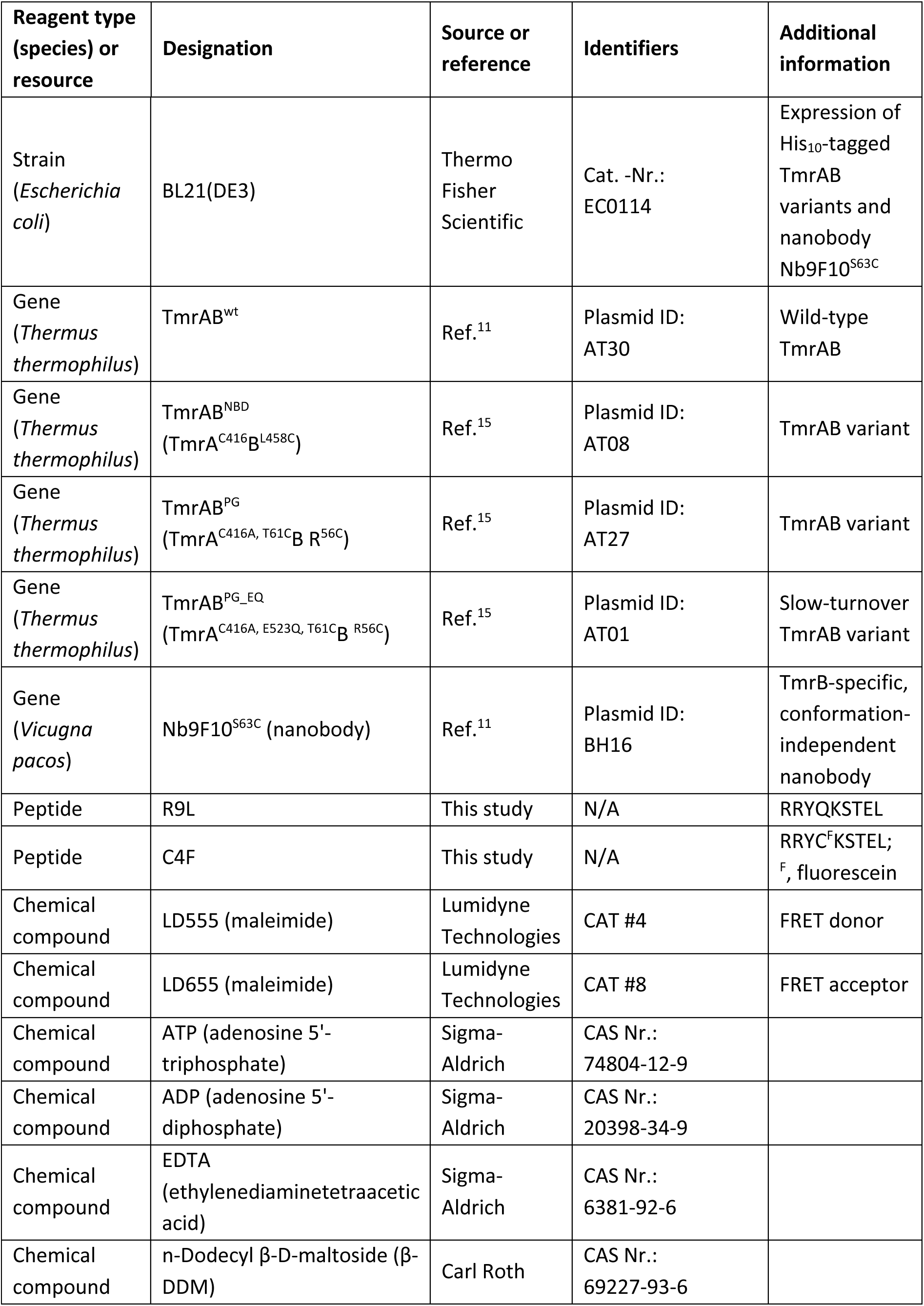
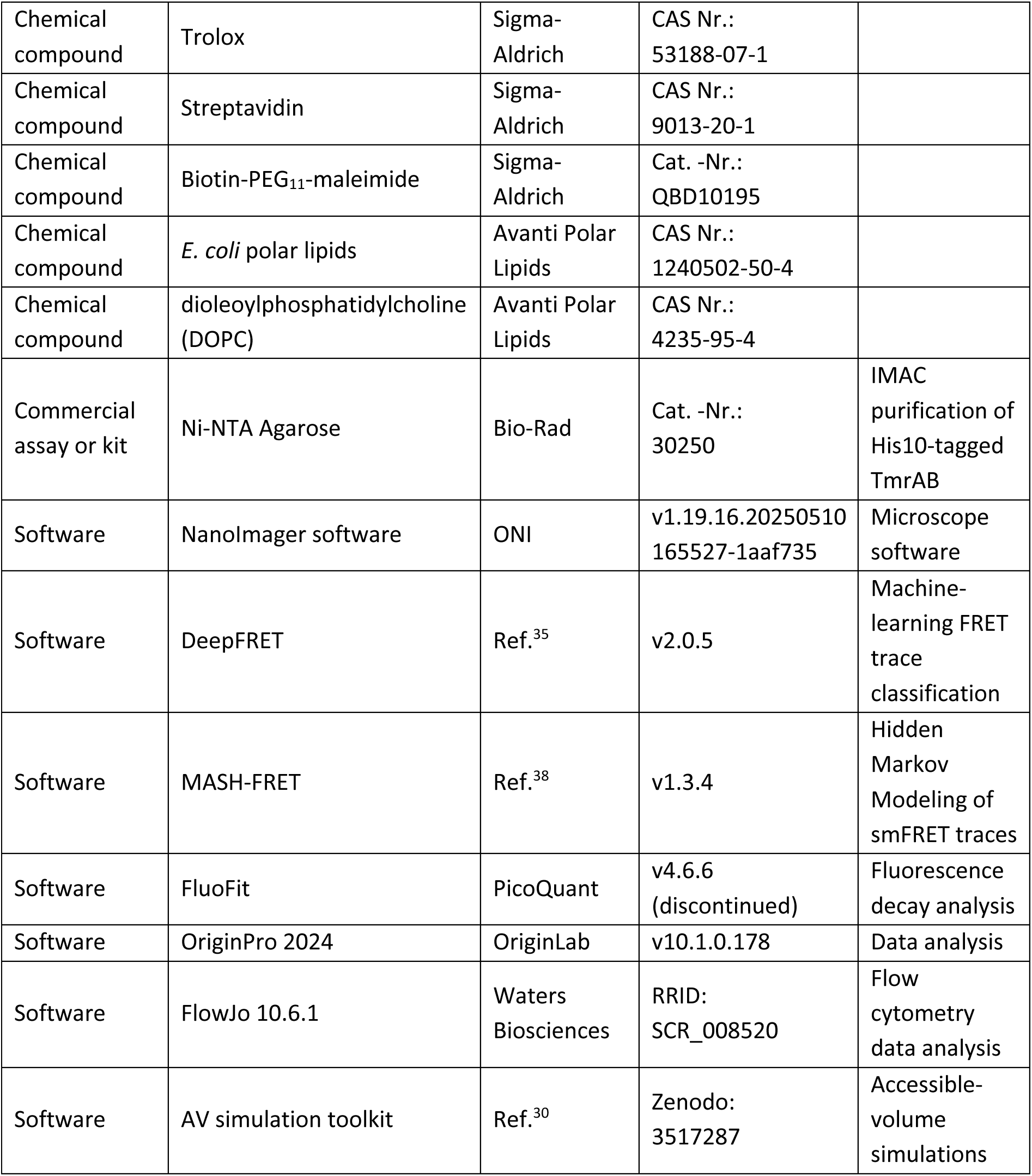

### Expression, purification, and labeling of TmrAB

His_10_-tagged TmrAB variants were expressed in *E. coli* BL21(DE3) (Thermo Fisher Scientific) as described previously^12^. Cells were grown in high-salt LB media (Carl Roth) supplemented with 100 μg ml^-1^ ampicillin (PAA Laboratories) at 37 °C. At an OD_600_ of 0.5, expression was induced with 1 mM isopropyl β-D-thiogalactopyranoside (IPTG; Carl Roth), and cultures were incubated for 3 h at 37 °C. Cells were harvested by centrifugation (4,500 × *g*, 4 °C, 15 min) and stored at –80 °C.

For purification, cell pellets were resuspended in lysis buffer (20 mM HEPES-NaOH pH 7.5, 300 mM NaCl, 50 µg ml^-1^ lysozyme, 0.2 mM phenylmethylsulfonyl fluoride (PMSF)) and lysed by sonication. Cell debris was removed by centrifugation (18,000 × *g*, 4 °C, 35 min), and membranes were collected by ultracentrifugation (100,000 × *g,* 4 °C, 30 min). Membranes were solubilized for 2 h at 4 °C in purification buffer (20 mM HEPES-NaOH pH 7.5, 300 mM NaCl) containing 20 mM n-dodecyl β-D-maltoside (β-DDM; Carl Roth). After ultracentrifugation (100,000 × *g,* 30 min, 4 °C), the supernatant was incubated with Ni-NTA agarose (Bio-Rad) for 1 h at 4 °C. The resin was washed with 20 column volumes of wash buffer (20 mM HEPES-NaOH pH 7.5, 300 mM NaCl, 1 mM β-DDM) containing 50 mM imidazole, and TmrAB was eluted with elution buffer (20 mM HEPES-NaOH pH 7.5, 300 mM NaCl, 1 mM β-DDM, 300 mM imidazole).

For fluorophore labeling, TmrAB variants were conjugated via maleimide chemistry using LD555 and LD655 (Lumidyne Technologies). Labeling was carried out at a 1:10:10 molar ratio of protein to each dye in elution buffer for 3 h at 4 °C. Excess dye was quenched with 2 mM β-mercaptoethanol (Sigma-Aldrich), and the labeled protein was buffer-exchanged into size-exclusion chromatography (SEC) buffer (20 mM HEPES-NaOH pH 7.5, 150 mM NaCl, 1 mM β-DDM) using a Zeba Spin Desalting Column (Thermo Fisher Scientific). Unreacted fluorophores were removed by SEC on a Superdex 200 Increase 10/300 GL column (Cytiva). Labeling efficiency was determined by analytical SEC (Superdex 200 Increase 3.2/300; Cytiva) by monitoring absorbance at 280, 555, and 655 nm. To preserve sample integrity for smFRET measurements, TmrAB was purified and labeled within a single day, stored on ice, and imaged over the following two days.

### Time-correlated single-photon counting (TCSPC)

Fluorescence lifetime measurements were performed using a FluoTime 100 spectrometer (PicoQuant) equipped for time-correlated single-photon counting (TCSPC). Experiments were carried out on labeled TmrAB in SEC buffer (20 mM HEPES-NaOH pH 7.5, 150 mM NaCl, 1 mM β-DDM). LD555 and LD655 were excited at 510 nm and 610 nm, respectively. Emission was collected using a 620/60 nm bandpass filter for LD555 and a BG4 700 nm long-pass filter for LD655. Photon arrival times were accumulated until the TCSPC histogram reached a peak count of 50,000 photons. Fluorescence decay curves were analyzed by fitting mono- or bi-exponential decay models using FluoFit software (PicoQuant) and amplitude-weighted average of fluorescence lifetime was calculated.

### Nanobody production and purification

The nanobody Nb9F10^S63C^ was expressed and purified as described previously^11^. Briefly, Nb9F10^S63C^ was produced in *E. coli* BL21(DE3) cells grown in Terrific Broth (TB; Carl Roth) supplemented with 100 μg ml^-1^ ampicillin at 37 °C. At an OD_600_ of 0.6, expression was induced with 1 mM IPTG, followed by overnight incubation at 28 °C. Cells were harvested by centrifugation (4,500 × *g*, 4 °C, 15 min) and stored at –80 °C. For purification, cell pellets were resuspended in nanobody lysis buffer (25 mM HEPES-NaOH pH 7.4, 300 mM NaCl, 15 mM imidazole, 0.5 mM PMSF) and disrupted by sonication. Cell debris was removed by centrifugation (18,000 × *g*, 4 °C, 35 min), and the clarified lysate was applied to Ni-NTA agarose equilibrated in potassium phosphate (KP_i_) buffer (25 mM KP_i_ pH 6.5, 100 mM KCl, and 0.5 mM tris(2-carboxyethyl) phosphine (TCEP)). Bound nanobody was washed with 10 column volumes (CV) of KP_i_ buffer and eluted with 8 CV of elution buffer (25 mM KP_i_ pH 6.0, 20 mM KCl, 300 mM imidazole, 0.5 mM TCEP). Eluted fractions were pooled and further purified by cation exchange chromatography (CEX) on a HiTrap SP column (Cytiva) using a linear gradient from low-salt buffer (25 mM KP_i_ pH 6.0, 20 mM KCl, 0.5 mM TCEP) to high-salt buffer (25 mM KP_i_ pH 6.0, 500 mM KCl, 0.5 mM TCEP). The purified nanobody was concentrated and buffer-exchanged into nanobody SEC buffer (20 mM HEPES-NaOH pH 7.5, 150 mM NaCl) using Zeba spin desalting columns, followed by SEC (Superdex 200 Increase 10/300 GL; Cytiva). For site-specific conjugation, Nb9F10^S63C^ was incubated with a biotin-PEG_11_-maleimide linker (Sigma-Aldrich) at a 1.2:1 molar ratio of protein to linker in the presence of 0.5 mM TCEP for 2 h at 4 °C. Excess linker was removed by desalting on Zeba Spin Desalting Columns, followed by a final SEC step (Superdex 200 Increase 10/300 GL).

### SDS-PAGE

The purity of TmrAB samples was assessed by SDS-PAGE. Resolving gels (10%) were prepared using 10% (w/v) acrylamide, 0.5 M Tris-HCl (pH 8.8), 0.13% (w/v) SDS, 0.05% (w/v) ammonium persulphate (APS), and 0.25% (v/v) *N,N,N’,N’-*tetramethylethylenediamine (TEMED). Stacking gels contained 4.3% (w/v) acrylamide, 0.5 M Tris/HCl (pH 6.8), 0.09% (w/v) SDS, 0.09% (w/v) APS, and 0.33% (v/v) TEMED. Gels were used immediately or stored at 4 °C for up to 4 weeks. Protein samples were mixed with 4× SDS loading buffer containing dithiothreitol (DTT; Sigma-Aldrich) and heated at 90 °C for 5 min before loading. Electrophoresis was performed at a constant voltage of 120 V using 1× SDS running buffer (25 mM Tris-HCl pH 8.8, 192 mM glycine, 0.1% SDS). Proteins were visualized by staining with InstantBlue^TM^ Protein Stain (Abcam) for 1 h at room temperature with gentle agitation and imaged using a Fusion FX system (Vilber).

### ATPase activity assay

The ATPase activity of β-DDM-solubilized TmrAB^wt^ was quantified using a Malachite Green-based colorimetric assay as described previously^44^. Detergent-solubilized TmrAB (60 nM) was incubated in ATPase buffer (20 mM HEPES-NaOH pH 7.5, 150 mM NaCl, 2 mM MgCl_2_, 1 mM β-DDM) containing 3 mM ATP (Sigma-Aldrich) at 40 °C for 7 min. Autohydrolysis controls were prepared by incubation of ATP in ATPase buffer without protein. Reactions were quenched by adding 20 mM H_2_SO_4_, followed by incubation with 3 mM Malachite Green (Thermo Fisher Scientific), 0.2% (v/v) Tween20 (Carl Roth), and 1.5% (w/v) ammonium molybdate (Carl Roth) for 10 min at room temperature. The absorbance at 620 nm was recorded on a CLARIOstar v.5.20 R5 plate reader (BMG LABTECH).

### Ensemble FRET measurements

The ATP binding and FRET characteristics of selected TmrAB variants were assessed by ensemble FRET. Labeled TmrAB (100 nM) was incubated with increasing concentrations of ATP at 42 °C for 5 min. Donor-excited emission was recorded from 550–700 nm with an excitation wavelength of 520 nm using a CLARIOstar v.5.20 R5 plate reader (BMG LABTECH). Acceptor emission intensities at 675 nm were plotted against ATP concentration and fitted with a hyperbolic function to determine the apparent dissociation constant (*K*_d, ATP_) for each variant.

### Reconstitution of TmrAB

TmrAB variants were reconstituted into proteoliposomes composed of *E. coli* polar lipids and dioleoylphosphatidylcholine (DOPC) at a molar ratio of 7:3. A protein-to-lipid ratio of 1:20 (w/w) was used, corresponding to approximately 50 TmrAB complexes per liposome. Extruded large unilamellar vesicles (LUVs) were destabilized by incubation with 0.3% (v/v) Triton X-100 for 30 min. TmrAB was then added to the detergent-destabilized liposomes and incubated for 30 min at 8 °C under gentle rotation. Detergent was removed gradually using sequential addition of polystyrene Bio-Beads SM-2 (Bio-Rad): two incubations at 40 mg ml^-1^ (1 h and overnight), followed by two incubations at 80 mg ml^-1^ (1 h each). Proteoliposomes were subsequently harvested by ultracentrifugation (100,000 × g, 30 min, 4 °C) and resuspended in the appropriate buffer.

### Single liposome transport assay

Single liposome transport assay was performed as described previously^12,13^. Wild-type TmrAB or LD555/LD655-labeled TmrAB^PG^ (0.6 µM each), reconstituted into liposomes (∼50 TmrAB per liposome), was mixed with fluorescently labeled peptide RRYC^F^KSTEL (30 µM C4F; ^F^, fluorescein), 3 mM ATP or ADP, and 5 mM MgCl_2_ in liposome buffer (25 mM HEPES-NaOH pH 7.5, 150 mM NaCl, 5% (v/v) glycerol). Where indicated, nanobody Nb9F10^S63C^ (0.6 µM) was added. Samples were incubated for 5 min at 40 °C (unless specified otherwise), followed by addition of ice-cold ethylenediaminetetraacetic acid (EDTA; 10 mM). Proteoliposomes were then washed twice by centrifugation (270,000 x g, 30 min) to remove non-transported peptide. Mean fluorescence intensities (MFIs) of translocated substrates were quantified by flow cytometry using a FACSMelody (Waters Biosciences) and analyzed in FlowJo 10.6.1 (Waters Biosciences). The gating strategy used to determine fluorescence intensities of single liposomes from raw flow cytometry data consisted of three steps: First, forward scatter area versus side scatter area (FSC-A vs SSC-A) was used to separate populations based on size and granularity. Second, forward scatter height versus forward scatter area (FSC-H vs FSC-A) was used to exclude aggregates and retain singlet events. Finally, fluorescence intensities of the gated singlet population were quantified at the wavelength of interest (527 nm), and the resulting values were used to calculate mean fluorescence intensity (MFI) of the proteoliposome population.

### Solid-phase peptide synthesis

The peptides used in this study were synthesized and labeled as described previously^14,45^. The 9-mer peptides RRYQKSTEL (R9L) and RRYC^F^KSTEL (C4F; ^F^, fluorescein) were synthesized on preloaded Fmoc-L-Leu resin using a Liberty microwave-assisted peptide synthesizer (CEM) according to a standard protocol (54 W, 3 min, 75 °C). Each coupling step was performed twice using 0.2 M fluorenylmethoxycarbonyl (Fmoc)-protected amino acids, 0.5 M O-benzotriazole-N,N,N’,N’-tetramethyluronium hexafluoro-phosphate (HBTU), and 1-hydroxybenzotriazole hydrate (HOBt H_2_O). Fmoc deprotection was carried out with 20% (v/v) piperidine and 0.1 M HOBt H_2_O in dimethylformamide (DMF). Peptides were cleaved from the resin using a cleavage cocktail containing 92.5% (v/v) trifluoroacetic acid (TFA), 2.5% (v/v) H_2_O, 4.5% (v/v) thioanisole, and 0.5% (v/v) 1,2-ethanedithiol (EDT) for 1.5 h at room temperature. Cleaved peptides were precipitated in ice-cold diethyl ether (Et_2_O), pelleted, dissolved in tert-butanol (*t*BuOH)/water (4:1, v/v), and lyophilized. Peptides were purified by reverse-phase (RP) C18 HPLC (PerfectSil C18 column; MZ-Analysentechnik) using buffer A (Milli-Q water containing 0.05% (v/v) TFA) and buffer B (acetonitrile containing 0.05% (v/v) TFA). Peptide identity and purity were confirmed by LC-MS. For fluorescence labeling, the peptide RRYC^F^KSTEL was dissolved in 33% (v/v) DMF in PBS (10 mM Na_2_HPO_4_, 1.8 mM KH_2_PO_4_, 137 mM NaCl, 2.7 mM KCl, pH 7.4) and incubated for 1 h at room temperature with a 1.3-fold molar excess of 5-iodoacetamido fluorescein (Merck).

### Functionalization of glass slides for single-molecule FRET analysis

Glass coverslips used for TmrAB immobilization in smFRET experiments were functionalized by PEGylation as described previously^46^. Coverslips (Carl Roth) were cleaned by sequential sonication in Milli-Q water and analytical-grade acetone (>99.9%; VWR International), followed by oxygen plasma treatment (0.3 mbar, 80% power, 15 min) using a Zepto plasma cleaner (Diener) and a 10 min incubation in methanol (Avantor, Gliwice, PL). Coverslips were then silanized by incubation for 30 min in a solution of 100 ml methanol, 5 ml acetic acid, and 3 ml 3-aminopropyltrimethoxysilane (APTES; Tokyo Chemical Industry). After silanization, slides were rinsed four times with methanol and dried under a nitrogen stream. Surface functionalization was achieved using a mixture of biotinylated-PEG (4 mol%) and nonbiotinylated-PEG (96 mol%; Rapp Polymer). The PEG solution was sandwiched between two coverslips and incubated overnight in a humidity chamber. Coverslips were then rinsed thoroughly with Milli-Q water and dried under nitrogen. To enhance passivation, a second PEGylation step was performed using 25 mM CH_3_-PEG-NHS (333 Da; Thermo Fisher Scientific) under the same conditions. Finally, slides were rinsed with Milli-Q water, dried under nitrogen, and stored at –20 °C under argon until use.

### Single-molecule FRET imaging

Single-molecule FRET (smFRET) experiments were performed using a flow chamber system (Ibidi). Chambers were assembled by placing a biotin-PEG-functionalized glass slide onto a μ-Slide I Luer Family flow channel (Ibidi), with the functionalized surface facing inward. All buffers and Milli-Q water were filtered through 0.2 μm filters (Sigma-Aldrich). Chambers were washed with 1 ml Milli-Q water and incubated with 0.2 mg ml^-1^ streptavidin (Sigma-Aldrich) at 4 °C for 30 min to allow binding to the biotin-PEG surface. Unbound streptavidin was removed by washing with 1 ml Milli-Q water. The surface was then treated with 0.3 mg ml^-1^ biotinylated-PEG_11_-Nb9F10^S63C^ at 4 °C for 45 min, followed by flushing with 2 ml SEC buffer (20 mM HEPES-NaOH pH 7.5, 150 mM NaCl, 1 mM β-DDM). Detergent-solubilized, fluorophore-labeled TmrAB (100 nM) was added and incubated at 4 °C for 1 h. Unbound protein was removed by five washes with 1 ml TmrAB-SEC buffer. Chambers were equilibrated with 1 ml of imaging buffer containing 25 mM HEPES-NaOH (pH 7.5), 150 mM NaCl, 3 mM MgCl_2_, 50 mM glucose, 5 mM Trolox, 7.5 U ml^-1^ pyranose oxidase, and 1 kU ml^-1^ catalase, supplemented with the desired ATP concentration. For EDTA trapping experiments, MgCl_2_ was omitted and replaced with 3 mM ethylenediaminetetraacetic acid (EDTA; Sigma-Aldrich). smFRET data were acquired at 40 °C using alternating laser excitation (ALEX) on a total internal reflection fluorescence (TIRF) microscope (NanoImager S, ONI, Oxford, UK). Typically, 600 frames were recorded per region of interest (ROI) with 100 ms exposure time. Laser powers were 0.8 mW cm^-2^ (532 nm) and 0.9 mW cm^-2^ (640 nm). Data were recorded in 1-min intervals, except for TmrAB^NBD^ variant in apo and 3 mM ATP conditions, where 3-min intervals were used to confirm that conformational transitions do not occur on timescales longer than one minute due to the reduced temperature.

### Single-molecule FRET data analysis

Single-molecule FRET (smFRET) measurements were performed using alternating laser excitation (ALEX), allowing assignment of detected photons based on both excitation and emission wavelengths. Photon counts for each molecule were extracted using NanoImager software (ONI NanoImager, Development Build) and classified into three detection channels: donor excitation with donor emission 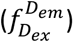, donor excitation with acceptor emission 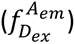, and acceptor excitation with acceptor emission 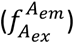. Traces were analyzed with DeepFRET^35^ and manually curated. To minimize bias, a second researcher independently curated traces from both ATP-free and 3 mM ATP samples, yielding 98% overlap between curations. FRET efficiency (*E*) and stoichiometry (*S*) were calculated as:

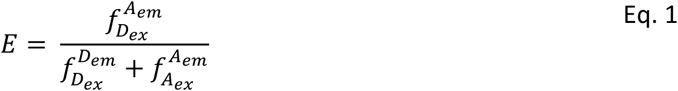

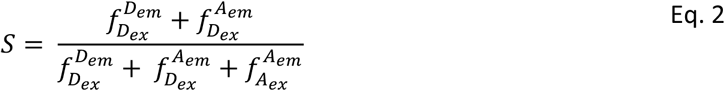

Population analysis was performed by constructing one-dimensional histograms of FRET efficiency (*E*) and stoichiometry (*S*) using OriginPro 2024 (OriginLab). Histograms were fitted with two Gaussian distributions corresponding to the ATP-free state (defined from apo samples) and the ATP-bound state (defined by a two-component fit at saturating ATP). Hidden Markov Modeling (HMM) of individual traces was performed using MASH-FRET^38^ to distinguish dynamic from static molecules within each sample.

#### ATP-bound dwell times and distribution of conformational states

The ATP-bound dwell time (*τ*_d_) was estimated as:

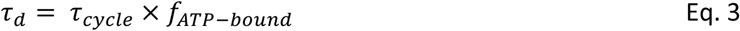

where *f*_ATP-bound_ is the fraction of ATP-bound molecules derived from Gaussian fits of the FRET efficiency histograms (**Fig. 3**). This approach assumes (i) ATP hydrolysis occurs exclusively at the canonical NBS, with negligible contribution from the noncanonical site^9,11,15^, and (ii) the majority of molecules are catalytically competent and continuously cycling.

The distribution of conformational states within ATP-bound FRET population (*E* = 0.86) of TmrAB^PG^ under turnover conditions (3 mM ATP) was determined using a two-state model:

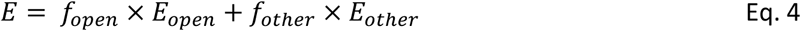

where *f*_open_ is the fraction of OF^open^ state (*E*_open_ = 0.63) readily resolved under trapping conditions (**Fig. 4**), and *f*_other_ represents the combined fraction of PG-closed states (OF^occluded^/UR^asym^/UR^asym^*, *E*_other_ = 0.97).

## Author contributions

M.P. prepared all TmrAB samples and carried out the experiments for this study. M.P. performed data analysis. Curated traces were independently checked by C.N. to avoid human bias. M.P. and C.N. prepared functionalized glass slides for single-molecule FRET. M.P. and R.T. wrote the manuscript. R.T. conceived and supervised the work.

## Acknowledgements

This work was supported by the European Research Council (ERC Advanced Grant 101141396 to R.T.), the German Research Foundation via the Collaborative Research Center CRC 1507/P18 to R.T. and the Research Training Group (GRK 1986/B4.7 to R.T.). We thank Jan F.M. Stuke and Jonas Göhmann for support in automating trace extraction from ONI NanoImager software, Dr. David Glück for guidance on lifetime measurements, and Tobias Nocker for preparing nanobodies used in the study. We are also grateful to the Wachtveitl lab (Goethe University Frankfurt) for access to their FluoTime 100 spectrometer (PicoQuant). Finally, we thank Dr. Rupert Abele, Dr. David Glück, Dr. Simon Trowitzsch, Inga Nold, and Andrea Pott for helpful comments and proofreading of the manuscript.

## Data and materials availability

All data are available in the main text or the supplementary materials. All other data are available from the corresponding author upon reasonable request. Source data are provided with this paper: DOI: https://doi.org/10.25716/gude.16a1-m6pe

## Competing Interest

The authors declare no competing interest.

---

**Figure 1-Figure supplement 1.**
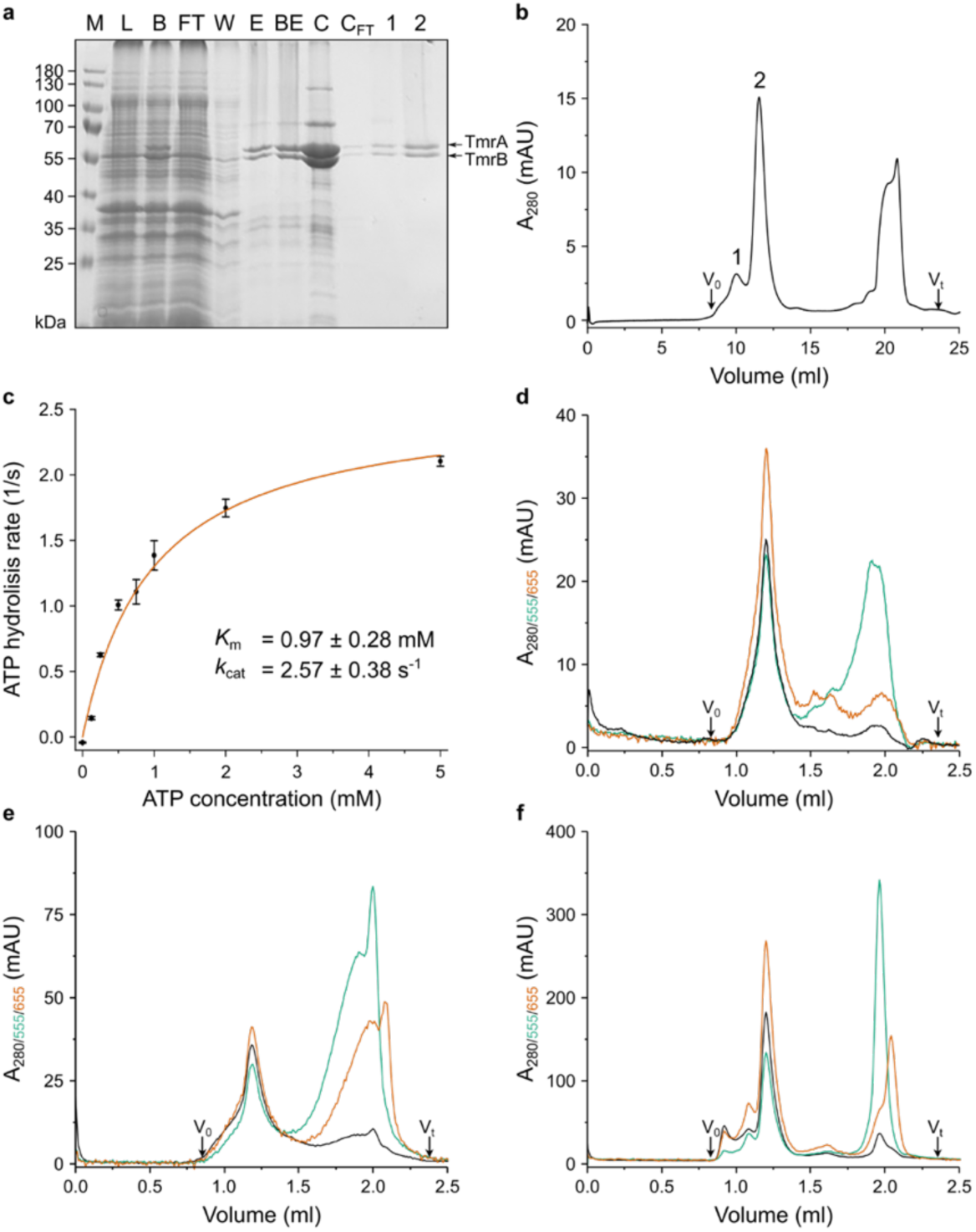
Quality of TmrAB purification and fluorophore labeling. **a**, SDS-PAGE analysis (10%, reducing gel, Coomassie staining) of successive purification steps: M, molecular weight marker; L, cell lysate; B, Ni-NTA beads after incubation with lysate; FT, flow-through; W, wash; E, eluted TmrAB; BE, buffer-exchanged sample; C, concentrated protein; CFT, concentrator flow-through; 1 and 2, first and second peaks eluted from size-exclusion chromatography (SEC). Only the second peak was used for subsequent FRET experiments. **b**, SEC (Superdex 200 increase 10/300 GL) showing monodisperse labeled TmrAB and efficient removal of free fluorophores. A representative chromatogram of TmrAB^PG^ is shown. **c**, ATP hydrolysis activity of purified wild-type TmrAB (60 nM TmrAB^wt^) measured at 40 °C for 7 min using the Malachite Green assay. Released inorganic phosphate (P_i_) was quantified and data fitted to Michaelis-Menten kinetics, yielding *K*_m_ = 0.97 ± 0.28 mM and *k*_cat_ = 2.57 ± 0.38 s^-1^. **d**–**f**, Analytical SEC (Superdex 200 increase 3.2/300) used to determine fluorophore labeling efficiencies of (**d**) TmrAB^NBD^, (**e**) TmrAB^PG^, and (**f**) TmrAB^PG_EQ^. LD555 and LD655 labeling efficiencies were ∼55 and ∼53% (TmrAB^NBD^), ∼43 and ∼52% (TmrAB^PG^), and ∼42% and ∼52% (TmrAB^PG_EQ^), respectively. Total cysteine occupancy exceeds 90% in all variants, reflecting near-complete dual-labeling with equimolar fluorophores.

**Figure 1-Figure supplement 2.**
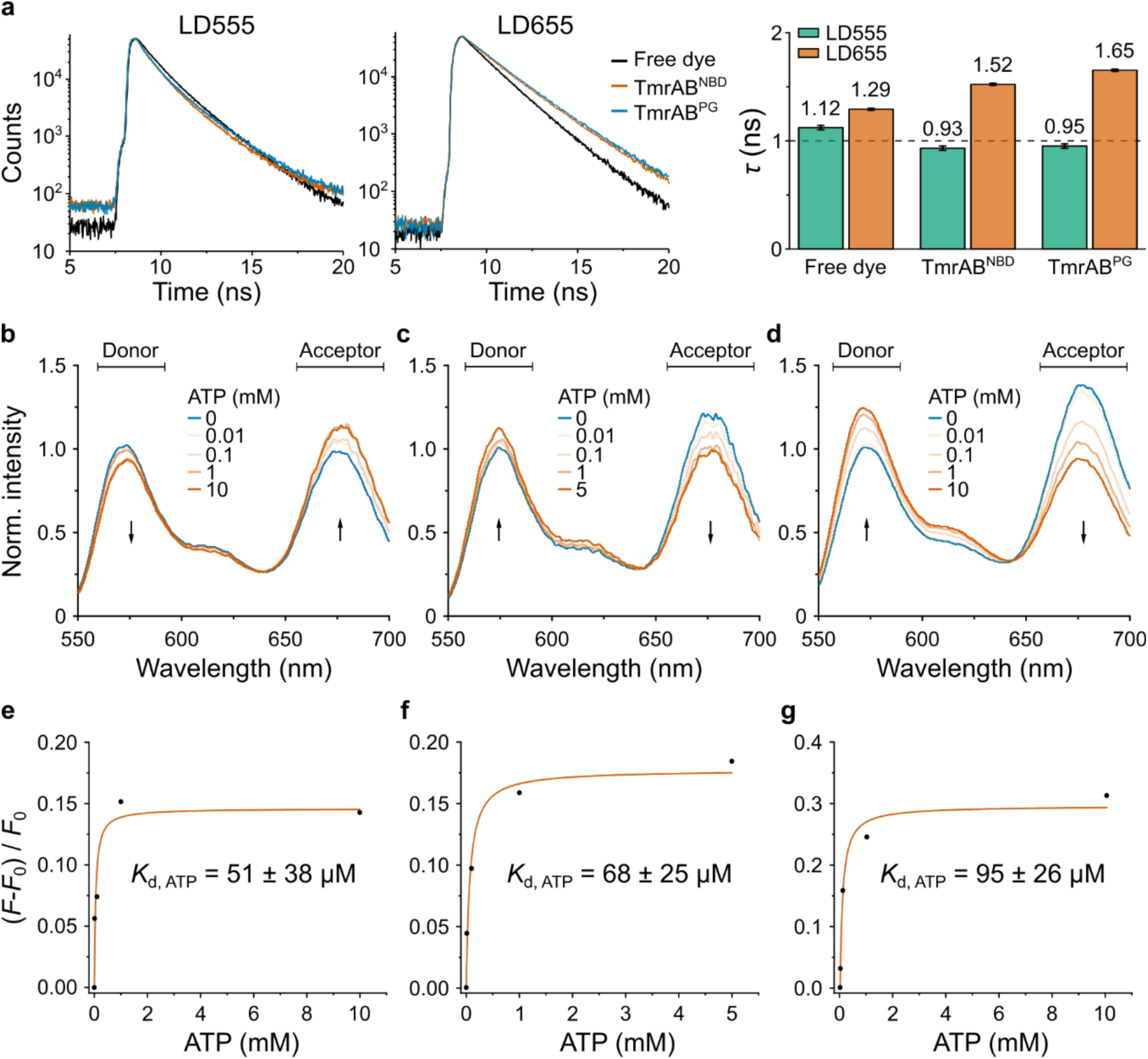
FRET properties of labeled TmrAB variants. **a**, Time-correlated single-photon counting histograms of LD555 (left) and LD655 (middle) measured for free dye in buffer (black), LD555/LD655-labeled TmrAB^NBD^ (orange), and LD555/LD655-labeled TmrAB^PG^ (blue). Amplitude-weighted average fluorescence lifetimes (*τ*) are summarized in the histogram (right), confirming sufficient fluorophore mobility for reliable FRET measurements. **b–d**, Donor-excited ensemble emission spectra (550–700 nm, excitation 520 nm) of stochastic LD555/LD655-labeled (**b**) TmrAB^NBD^, (**c**) TmrAB^PG^, and (**d**) the slow-turnover variant TmrAB^PG_EQ^ measured at increasing ATP concentrations. Spectra were normalized to donor intensity in the apo state. ATP-dependent donor quenching and acceptor sensitization demonstrate that all variants retain FRET capability. **e–g**, Fractional fluorescence changes, (*F*-*F*₀)/*F*₀, plotted as a function of ATP concentration for (**e**) TmrAB^NBD^, (**f**) TmrAB^PG^, and (**g**) TmrAB^PG_EQ^, where *F* is the acceptor emission intensity and *F*₀ the intensity in the apo state. Data were fitted with a hyperbolic (Langmuir-type) binding model to determine apparent *K*_d, ATP_ values, which are consistent with ensemble FRET measurements of ATP binding.

**Figure 1-Figure supplement 3.**
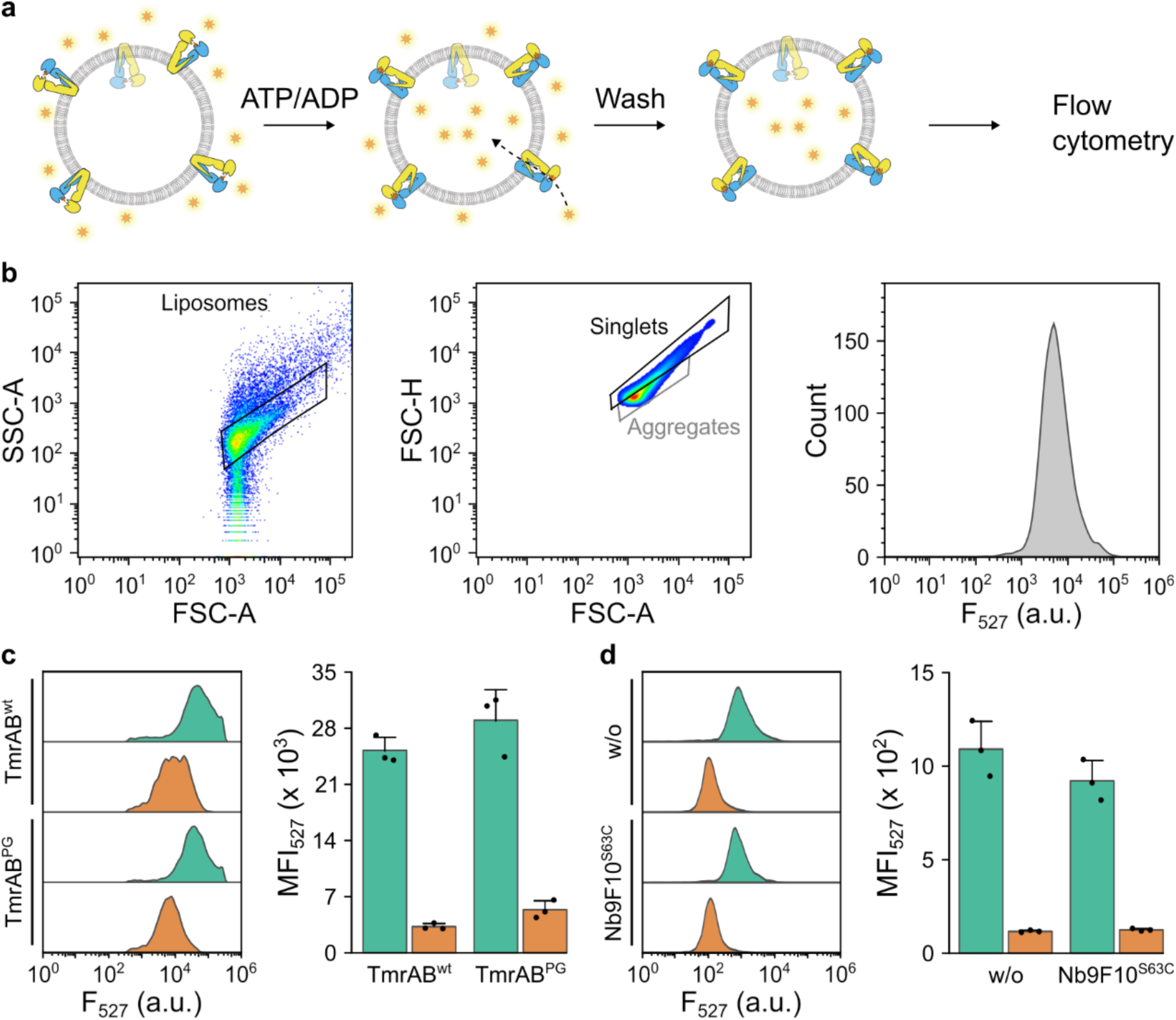
Effect of fluorophore labeling and nanobody binding on TmrAB transport activity. Single-liposome transport assays were performed by flow cytometry using mean fluorescence intensity at 527 nm (MFI_527_) as a readout of peptide transport. **a**, TmrAB (0.6 µM) reconstituted into liposomes (∼50 transporters per liposome) was incubated with C4F peptide (30 µM; RRYC^F^KSTEL, ^F^, fluorescein) in the presence of 3 mM ATP (green) or 3 mM ADP (orange) at 40 °C for 5 min. After extensive washing, proteoliposomes were analyzed by flow cytometry. **b**, Flow-cytometry gating strategy. Forward scatter area (FSC-A) versus side scatter area (SSC-A; left) was used to identify homogeneous proteoliposomes, forward scatter height (FSC-H) versus FSC-A (middle) to select singlets, and fluorescence intensity at 527 nm (F_527_; right) to determine the mean fluorescence intensity (MFI) values of the gated population. **c**, Transport activity of unlabeled wild-type TmrAB (TmrAB^WT^) compared with LD555/LD655-labeled TmrAB^PG^. **d**, Effect of nanobody binding on TmrAB^wt^ transport activity. Assays were performed in the absence or presence of Nb9F10^S63C^ at a 1:1 molar ratio (0.6 µM each). Fluorophore labeling at the periplasmic gate and nanobody binding did not measurably affect TmrAB transport activity.

**Figure 2-Figure supplement 1.**
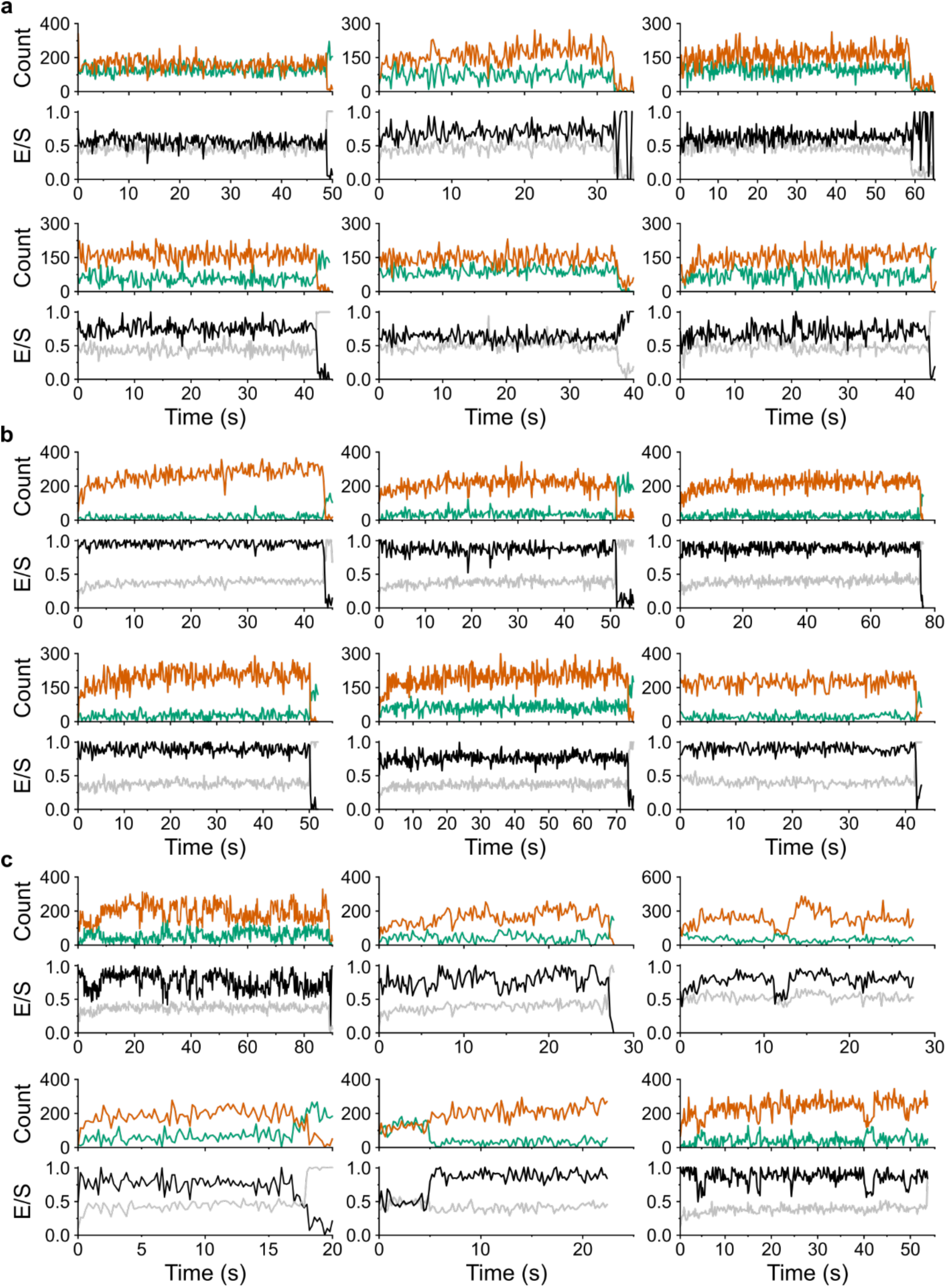
Representative smFRET traces of TmrAB^NBD^. Representative single-molecule FRET (smFRET) traces of TmrAB^NBD^ recorded (**a**) in the absence of ATP and (**b, c**) in the presence of 3 mM ATP. Hidden Markov modeling (HMM) classified ATP-bound traces as (**b**) static or (**c**) dynamic based on the absence or presence of transitions between ATP-free and ATP-bound conformational states. Donor fluorescence upon donor excitation is shown in green, acceptor fluorescence upon donor excitation in orange, FRET efficiency (*E*) in black, and stoichiometry (*S*) in grey.

**Figure 2-Figure supplement 2.**
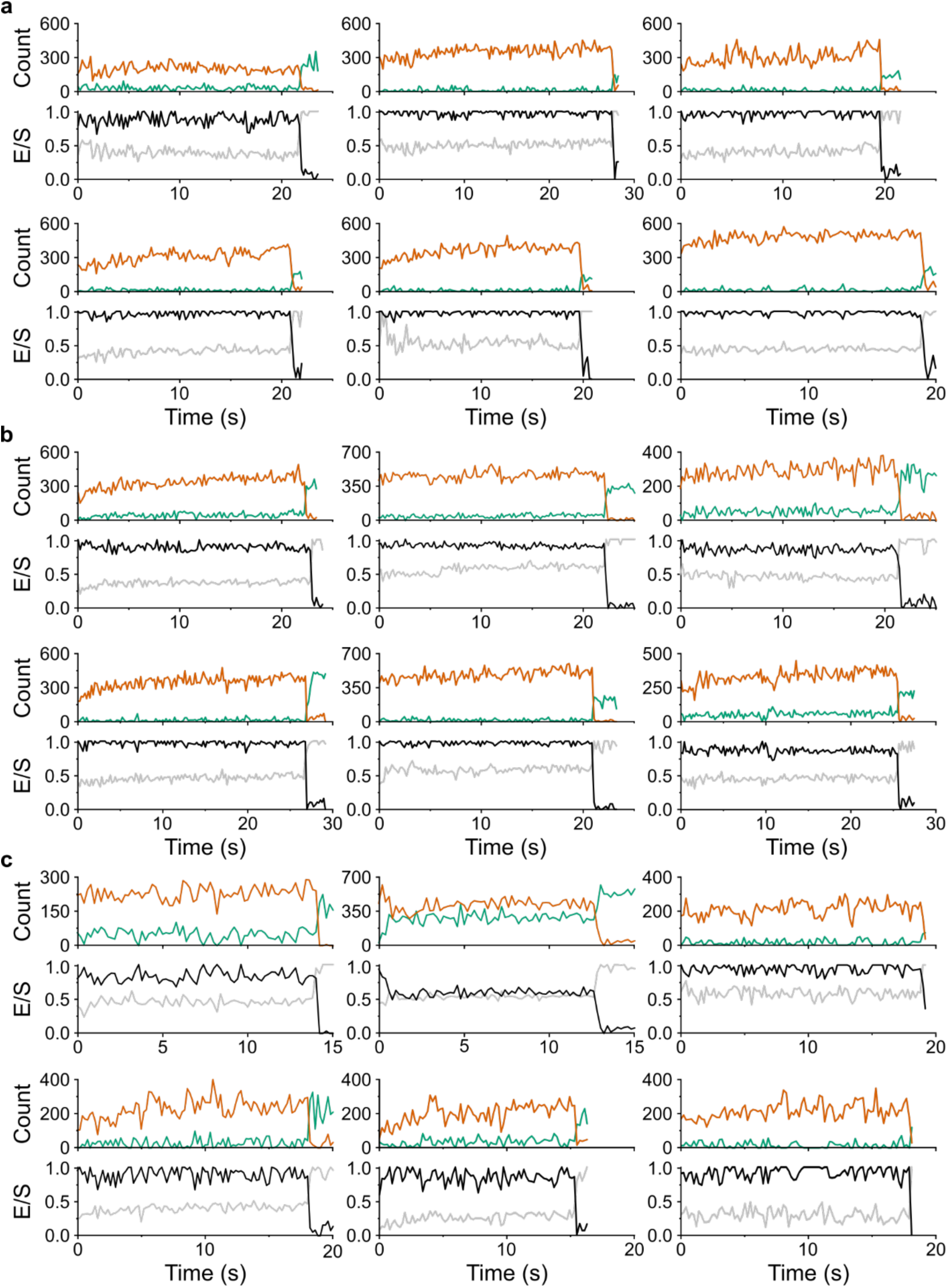
Representative smFRET traces of TmrAB^PG^. Representative single-molecule FRET (smFRET) traces of TmrAB^PG^ recorded (**a**) in the absence of ATP and (**b, c**) in the presence of 3 mM ATP. Hidden Markov modeling (HMM) classified ATP-bound traces as (**b**) static or (**c**) dynamic based on the absence or presence of transitions between ATP-free and ATP-bound conformational states. Donor fluorescence upon donor excitation is shown in green, acceptor fluorescence upon donor excitation in orange, FRET efficiency (*E*) in black, and stoichiometry (*S*) in grey.

## Notes

### Competing Interest Statement

The authors have declared no competing interest.

### Summary of Updates

We have revised the final section of the Results to better reflect the scope indicated by the heading.

## References

1. Davidson AL, Dassa E, Orelle C, Chen J. Structure, function, and evolution of bacterial ATP-binding cassette systems. Microbiol Mol Biol Rev 72, 317–364 (2008).

2. Rees DC, Johnson E, Lewinson O. ABC transporters: the power to change. Nat Rev Mol Cell Biol 10, 218–227 (2009).

3. Thomas C, Tampé R. Structural and mechanistic principles of ABC transporters. Annu Rev Biochem 89, 605–636 (2020).

4. Locher KP. Mechanistic diversity in ATP-binding cassette (ABC) transporters. Nat Struct Mol Biol 23, 487–493 (2016).

5. Thomas C, et al. Structural and functional diversity calls for a new classification of ABC transporters. FEBS Lett 594, 3767–3775 (2020).

6. Robey RW, Pluchino KM, Hall MD, Fojo AT, Bates SE, Gottesman MM. Revisiting the role of ABC transporters in multidrug-resistant cancer. Nat Rev Cancer 18, 452–464 (2018).

7. Abele R, Tampé R. The ABCs of immunology: structure and function of TAP, the transporter associated with antigen processing. Physiology (Bethesda) 19, 216–224 (2004).

8. Kim J, et al. Subnanometre-resolution electron cryomicroscopy structure of a heterodimeric ABC exporter. Nature 517, 396–400 (2015).

9. Zutz A, et al. Asymmetric ATP hydrolysis cycle of the heterodimeric multidrug ABC transport complex TmrAB from *Thermus thermophilus*. J Biol Chem 286, 7104–7115 (2011).

10. Nöll A, et al. Crystal structure and mechanistic basis of a functional homolog of the antigen transporter TAP. Proc Natl Acad Sci U S A 114, E438–E447 (2017).

11. Hofmann S, et al. Conformation space of a heterodimeric ABC exporter under turnover conditions. Nature 571, 580–583 (2019).

12. Stefan E, Hofmann S, Tampé R. A single power stroke by ATP binding drives substrate translocation in a heterodimeric ABC transporter. eLife 9, e55943 (2020).

13. Stefan E, et al. De novo macrocyclic peptides dissect energy coupling of a heterodimeric ABC transporter by multimode allosteric inhibition. eLife 10, e67732 (2021).

14. Nocker C, Pečak M, Nocker T, Fahim A, Sušac L, Tampé R. Single-molecule dynamics reveal ATP binding alone powers substrate translocation by an ABC transporter. Nat Commun 17, (2026).

15. Barth K, Rudolph M, Diederichs T, Prisner TF, Tampé R, Joseph B. Thermodynamic basis for conformational coupling in an ATP-binding cassette exporter. J Phys Chem Lett 11, 7946–7953 (2020).

16. Barth K, Hank S, Spindler PE, Prisner TF, Tampé R, Joseph B. Conformational coupling and trans-inhibition in the human antigen transporter ortholog TmrAB resolved with dipolar EPR spectroscopy. J Am Chem Soc 140, 4527–4533 (2018).

17. Agam G, et al. Reliability and accuracy of single-molecule FRET studies for characterization of structural dynamics and distances in proteins. Nat Methods 20, 523–535 (2023).

18. Hellenkamp B, et al. Precision and accuracy of single-molecule FRET measurements-a multi-laboratory benchmark study. Nat Methods 15, 669–676 (2018).

19. Sasmal DK, Pulido LE, Kasal S, Huang J. Single-molecule fluorescence resonance energy transfer in molecular biology. Nanoscale 8, 19928–19944 (2016).

20. Bartels K, Lasitza-Male T, Hofmann H, Löw C. Single-molecule FRET of membrane transport proteins. ChemBioChem 22, 2657–2671 (2021).

21. Lerner E, et al. FRET-based dynamic structural biology: Challenges, perspectives and an appeal for open-science practices. eLife 10, e60416 (2021).

22. Nettels D, Galvanetto N, Ivanović MT, Nüesch M, Yang T, Schuler B. Single-molecule FRET for probing nanoscale biomolecular dynamics. Nat Rev Phys 6, 587–605 (2024).

23. Wang L, Johnson ZL, Wasserman MR, Levring J, Chen J, Liu S. Characterization of the kinetic cycle of an ABC transporter by single-molecule and cryo-EM analyses. eLife 9, e56451 (2020).

24. Levring J, Terry DS, Kilic Z, Fitzgerald G, Blanchard SC, Chen J. CFTR function, pathology and pharmacology at single-molecule resolution. Nature 616, 606–614 (2023).

25. Husada F, et al. Conformational dynamics of the ABC transporter McjD seen by single-molecule FRET. EMBO J 37, e100056 (2018).

26. Grossmann N, Vakkasoglu AS, Hulpke S, Abele R, Gaudet R, Tampé R. Mechanistic determinants of the directionality and energetics of active export by a heterodimeric ABC transporter. Nat Commun 5, 5419 (2014).

27. Sun P, et al. Substrate recognition diversity and transport dynamics of ABCC1. Nat Commun 16, 10499 (2025).

28. Altman RB, et al. Cyanine fluorophore derivatives with enhanced photostability. Nat Methods 9, 68–71 (2011).

29. Martin MI, et al. Leveraging Baird aromaticity for advancement of bioimaging applications. J Phys Org Chem 36, e4449 (2023).

30. Kalinin S, et al. A toolkit and benchmark study for FRET-restrained high-precision structural modeling. Nat Methods 9, 1218–1225 (2012).

31. Ha T, Tinnefeld P. Photophysics of fluorescent probes for single-molecule biophysics and super-resolution imaging. Annu Rev Phys Chem 63, 595–617 (2012).

32. Sindbert S, et al. Accurate distance determination of nucleic acids via Förster resonance energy transfer: implications of dye linker length and rigidity. J Am Chem Soc 133, 2463–2480 (2011).

33. Dale RE, Eisinger J, Blumberg WE. The orientational freedom of molecular probes. The orientation factor in intramolecular energy transfer. Biophys J 26, 161–193 (1979).

34. van der Meer BW. Kappa-squared: from nuisance to new sense. J Biotechnol 82, 181–196 (2002).

35. Thomsen J, et al. DeepFRET, a software for rapid and automated single-molecule FRET data classification using deep learning. eLIfe 9, e60404 (2020).

36. Hohlbein J, Craggs TD, Cordes T. Alternating-laser excitation: single-molecule FRET and beyond. Chem Soc Rev 43, 1156–1171 (2014).

37. Asher WB, et al. Single-molecule FRET imaging of GPCR dimers in living cells. Nat Methods 18, 397–405 (2021).

38. Hadzic M, Borner R, Konig SLB, Kowerko D, Sigel RKO. Reliable state identification and state transition detection in fluorescence intensity-based single-molecule Förster Resonance Energy-Transfer data. J Phys Chem B 122, 6134–6147 (2018).

39. Dimura M, Peulen TO, Hanke CA, Prakash A, Gohlke H, Seidel CA. Quantitative FRET studies and integrative modeling unravel the structure and dynamics of biomolecular systems. Curr Opin Struct Biol 40, 163–185 (2016).

40. Lam K, Tajkhorshid E. Membrane interactions of Cy3 and Cy5 fluorophores and their effects on membrane-protein dynamics. Biophys J 119, 24–34 (2020).

41. Grabenhorst L, Sturzenegger F, Hasler M, Schuler B, Tinnefeld P. Single-molecule FRET at 10 MHz count rates. J Am Chem Soc 146, 3539–3544 (2024).

42. Bonhomme L, et al. Triple labeling resolves a GPCR intermediate state by using three-color single molecule FRET. J Am Chem Soc 147, 17689–17700 (2025).

43. Yim SW, et al. Four-color alternating-laser excitation single-molecule fluorescence spectroscopy for next-generation biodetection assays. Clin Chem 58, 707–716 (2012).

44. Diederichs T, Tampé R. Single cell-like systems reveal active unidirectional and light-controlled transport by nanomachineries. ACS Nano 15, 6747–6755 (2021).

45. Löffler M, Frühschulz S, Rockel Z, Pečak M, Tampé R, Wieneke R. Antigen Delivery Controlled by an On-Demand Photorelease. Angew Chem Int Ed Engl 63, e202405035 (2024).

46. Chandradoss SD, Haagsma AC, Lee YK, Hwang JH, Nam JM, Joo C. Surface passivation for single-molecule protein studies. J Vis Exp, 50549 (2014).

47. Chaptal V, et al. Substrate-bound and substrate-free outward-facing structures of a multidrug ABC exporter. Sci Adv 8, eabg9215 (2022).

